# The acidic intrinsically disordered region of the inflammatory mediator HMGB1 mediates fuzzy interactions with chemokine CXCL12

**DOI:** 10.1101/2023.06.06.543836

**Authors:** Malisa Vittoria Mantonico, Federica De Leo, Giacomo Quilici, Liam Colley, Francesco De Marchis, Massimo Crippa, Tim Schulte, Chiara Zucchelli, Stefano Ricagno, Gabriele Giachin, Michela Ghitti, Marco Bianchi, Giovanna Musco

## Abstract

Chemokines engage in heterodimeric interactions to activate or dampen their cognate receptors in inflammatory conditions. The chemokine CXCL12 forms with the alarmin HMGB1 a patho-physiologically relevant heterocomplex (HMGB1●CXCL12), whose formation synergically promotes the inflammatory response elicited by the G-protein coupled receptor CXCR4. However, the molecular details of complex formation were still elusive. Through an integrative structural approach (NMR, AUC, ITC, MST, SAXS) we show that HMGB1●CXCL12 represents the first *fuzzy* chemokines heterocomplex reported so far. HMGB1 and CXCL12 form a dynamic equimolar assembly, rather than involving one HMGB1 and two CXCL12 molecules as previously assumed, with structured and unstructured HMGB1 regions recognizing the dimerization surface of CXCL12. We uncover an unexpected role of the acidic intrinsically disordered region (IDR) in heterocomplex formation and provide the first evidence that the acidic IDR facilitates the ternary HMGB1•CXCL12•CXCR4 interaction on the cell surface. Thus, the interaction of HMGB1 with CXCL12 diverges radically from the classical rigid heterophilic chemokine-chemokine dimerization. Simultaneous interference with the multiple interactions within HMGB1●CXCL12 complex formation might offer novel pharmacological strategies to inhibit its detrimental activity in inflammatory conditions.

## Introduction

Chemokines constitute a large family of signaling proteins that interact with cell surface G protein-coupled chemokine receptors (GPCRs). Through an intricate network of cross-talks with their receptors they regulate leukocyte activation and trafficking in physiological and pathological conditions^1, 2^. Chemokines are characterized by a conserved, three-stranded β-sheet/α-helix fold and exist in a monomer-multimer equilibrium. A shift towards one or the other form can either activate or dampen their cognate receptors, thus adding a further layer of complexity to the functional tuning of chemokine/receptor axis and of their downstream pathways^3–5^. Another sophisticated mechanism of chemokine fine-regulation is their ability to heterodimerize with other chemokines, resulting in synergic activation and dimerization of their cognate receptors^6–8^. This mechanism is particularly relevant in inflammatory conditions, where chemokine heterocomplexes activate chemokine receptors in the presence of low concentrations of chemokine-selective agonists that otherwise, without the synergy-inducing partner, would be inactive^8, 9^. Examples include the interaction of (i) CCL21 or CCL19 with CXCL13 to enhance leukocyte migration and activities through binding and activation of CCR7 at lower agonist concentrations^10^, (ii) CCL5 with CXCL4 to recruit monocytes and neutrophils^11^, and (iii) CXCL12 and CXCL9 with CXCR4 to attract lymphoma cells^12^. Such a *chemokine interactome* further enlarges the possibility of heteromeric interactions among chemokines, expanding the fine-regulation of signaling possibilities^11, 13^.

Intriguingly, chemokines can also form heterophilic interactions with some inflammatory mediators that are not structurally homologous to the classical CC-, CXC-, CX3C-, or XC- chemokines^5^. In this sense CXCL12 (chemokine (C-X-C motif) ligand 12) represents a paradigmatic example, as it is also able to bind to other proteins, such as galectins^14^ and the alarmin High Mobility Group Box 1 (HMGB1)^6, 15^. The interaction with Galectin 3 is immunoregulatory and attenuates CXCL12-stimulated signaling via CXCR4^14^. Conversely, binding of CXCL12 to HMGB1 synergistically enhances the CXCR4-dependent chemotactic response of monocytes and is involved in tissue regeneration^6, 15^. The 25 kDa HMGB1 protein comprises two L-shaped HMG tandem boxes (∼80 aa each connected by a flexible linker), referred to as BoxA and BoxB, and an acidic C-terminal intrinsically disordered region (IDR, 30 aa). HMGB1 is a Damage Associated Molecular Pattern (DAMP), which once released in the extracellular space alerts the host to stress, unscheduled cell death or microbial invasion, thus triggering inflammation and immune responses^16, 17^. HMGB1 is passively released by dead non-apoptotic cells and is actively released by severely stressed cells and by immune cells such as macrophages, natural killer cells, neutrophils and mature dendritic cells (reviewed in ^18^). It contains three cysteines (C22, C44, and C105), whose redox states, a disulfide (ds-HMGB1) and a fully reduced form (fr-HMGB1), determine how HMGB1 functions as a pro-inflammatory mediator^19^. On one hand, ds-HMGB1, with a disulphide bridge located on BoxA, binds to the Toll-like receptor 4/MD2 complex herewith promoting inflammatory response and cytokines activation^20^. Conversely, fr-HMGB1 plays a pivotal role in promoting the recruitment of inflammatory cells to injured tissues via heterocomplex formation with CXCL12 (HMGB1●CXCL12) and activation of CXCR4. This heterocomplex induces specific CXCR4 homodimer rearrangements, promotes CXCR4-mediated signaling, resulting in increased ERK activation and calcium rise induction^15^ and maintains CXCR4 on the plasma membrane in a β-arrestin 2 dependent manner^21^.

While Galectin 3 is structurally reminiscent of chemokines and interacts with CXCL12 exploiting in part the CXC type dimerization surface, composed by the beta-strand β1 and the alpha-helix α1^14^, the molecular details dictating HMGB1●CXCL12 intermolecular interactions are in part elusive, possibly because of the intrinsic dynamics of the system components. On one side CXCL12, as a typical chemokine, exists in a physiologically relevant monomer-dimer equilibrium^22^. On the other side, HMGB1 oscillates between a collapsed and open form, via intramolecular electrostatic interaction between the acidic IDR and the basic HMG boxes^23–25^, thus adding a further degree of complexity to the system^22^. Whether one or two molecules of CXCL12 bind to HMGB1 and whether the acidic IDR plays a role in heterocomplex formation and function are still open questions^22, 26^. Here, to address these issues, we have adopted a dissection and mutagenesis strategy, coupled to an integrative structural approach (NMR, AUC, ITC, MST, SAXS). Our results reveal that HMGB1 and CXCL12 associate via *fuzzy* interactions, forming an equimolar, highly dynamic heterocomplex. The acidic IDR plays a hitherto neglected but prominent role in complex formation *in vitro* and within a cellular environment. Indeed, the HMGB1●CXCL12 heterocomplex binds the CXCR4 receptor, and this binding is facilitated by the acidic IDR.

## Results

### The HMGB1 acidic IDR participates to the formation of the HMGB1●CXCL12 heterocomplex

The molecular details and the actual role of the single HMGB1 domains in the formation of the heterocomplex with CXCL12 are still elusive. Existing models, based on NMR titrations between the single HMG boxes and CXCL12, are based on the assumption that only the structured domains are the main actors in complex formation^26, 27^. Whether the acidic IDR of HMGB1 plays a role in complex formation has never been explored. We have therefore adopted a dissecting approach and performed comparative nuclear magnetic resonance (NMR) experiments titrating ^15^N CXCL12 with a synthetic peptide corresponding to the acidic IDR (Ac-pep), a tail-less HMGB1 construct, composed of the HMG tandem domain devoid of the acidic IDR (HMGB1-TL), and full-length HMGB1 (**Figure 1**).

**Figure 1:**
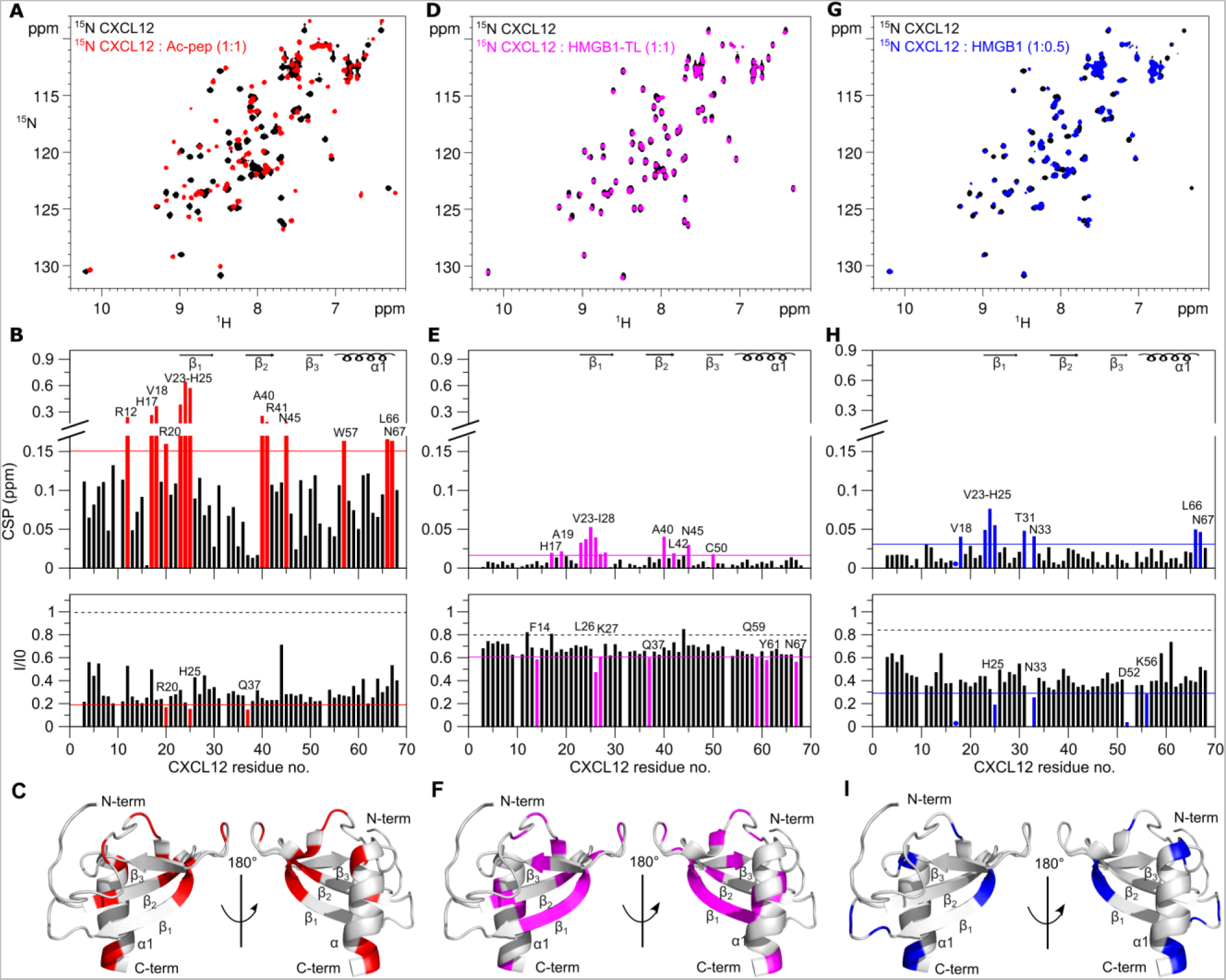
Ac-pep, HMGB1-TL and HMGB1 interact with CXCL12 dimerization surface. **(A)** Superposition of ^1^H-^15^N HSQC spectra of ^15^N CXCL12 (0.1 mM) without (black) and with (red) Ac-pep (1:1). **(B)** Bar graph showing residue-specific CSPs (upper panel) and intensities ratios (I/I_0_) (lower panel) of ^15^N-labeled CXCL12 (0.1 mM) upon addition of Ac-pep (1:1). Residues with CSP > Avg + σ_0_ (corrected standard deviation, red line) and with I/I_0_ < Avg - SD (standard deviation, red line) are labeled and **(C)** shown in red on CXCL12 (gray cartoon, pdb code:2kee). **(D)** Superposition of ^1^H-^15^N HSQC spectra of ^15^N CXCL12 (0.1 mM) without (black) and with (magenta) HMGB1-TL (1:1). **(E)** Bar graph showing residue-specific CSPs (upper panel) and intensities ratios (I/I_0_) (lower panel) of ^15^N-labeled CXCL12 (0.1 mM) upon addition of HMGB1-TL (1:1) of ^15^N-labeled CXCL12 (0.1 mM). Residues with CSP > Avg + σ_0_ (magenta line) with I/I_0_ < Avg - SD (magenta line) are labeled and **(F)** shown in magenta on CXCL12. **(G)** Superposition of ^1^H-^15^N HSQC spectra of ^15^N CXCL12 (0.1 mM) without (black) and with (blue) HMGB1 (1:0.5). **(H)** Bar graph showing residue-specific CSPs (top panel) and intensities ratios (I/I_0_) of ^15^N-labeled CXCL12 (0.1 mM) upon addition of HMGB1 (1:0.5). Residues with CSP > Avg + σ_0_ (blue line) and I/I_0_ < Avg - SD (blue line) are labeled and (**I**) shown in blue on CXCL12. In the bar-graphs α-helices and β-strands are schematically represented on the top, missing residues are prolines, dots indicate residues disappearing upon binding, the dashed black line indicates the expected peak intensity decrease due to the titration dilution effect.

Notably, addition of Ac-pep to ^15^N CXCL12 induced a dramatic change of the corresponding ^1^H-^15^N Heteronuclear Single Quantum Coherence (HSQC) spectrum, with substantial chemical shift perturbations (CSPs, in the 0.2-0.6 ppm range) and overall peak intensity reduction (**Figure 1A, B**). Peaks were severely broadened or disappeared beyond detection upon addition of sub-stoichiometric amounts of Ac-pep, but reappeared at 1:1 stoichiometry (**Supplementary Figure S1A)**. The dissociation constant was in the sub-micromolar range (Kd = 0.2 ± 0.1µM), as assessed by line-shape analysis using the software TITAN^28^ (**Supplementary Figure S1B**). CXCL12 residues with the highest CSPs were R12, H17, V18, R20, V23, K24, H25, A40, R41, N45, W57, L66, N67 (**Figure 1B**), with V23, K24, H25 (on β1) and L66, N67 (on α1) being located on the known CXCL12 homodimerization interface (**Figure 1C**).

In contrast to Ac-pep, addition of HMGB1-TL to ^15^N CXCL12 (**Figure 1D, E**) induced very small CSPs (in the 0.02-0.06 ppm range). As typically observed in NMR studies of chemokine heterocomplex formation, the interaction occurred in the intermediate exchange regime on the NMR chemical shift timescale^14^, with peak-intensity reduction upon binding. Residues mostly affected by the interaction with HMGB1-TL partially coincided with the ones affected by Ac-pep (H17, V23-H25, A40, N45, N67), suggesting that both the acidic IDR and the HMG tandem domain in part share the same interaction surface (**Figure 1F**). Similarly, titration of full-length HMGB1 induced significant CSPs (in the 0.05-0.1ppm range) on residues located on the β1 strand (V23-H25) and α-helix (L66, N67) (**Figure 1G-I**). Herein, we observed pronounced line broadening effects already at sub-stoichiometric concentrations (1:0.5) that hampered analysis at equimolar ratio, suggesting different binding dynamics and affinities for the full-length protein and its different constructs. Of note, comparison of the profiles of the CSPs of ^15^N CXCL12 upon addition of either Ac-pep, HMGB1-TL or HMGB1 were similar, with residues located on CXCL12 dimerization surface, showing the highest CSPs (**Figure 1C**). Importantly, the chemical shifts of these residues and their perturbations strongly depend on the CXCL12 monomer-dimer equilibrium^22^. Thus, to distinguish CSPs due to direct interactions with HMGB1 and constructs thereof from those related to potential changes in the CXCL12 oligomerization state, we repeated the titrations with a CXCL12 mutant locked in a monomeric state (CXCL12-LM)^29^. The spectral perturbations upon Ac-pep, HMGB1-TL and HMGB1 addition were similar to those observed with wild type CXCL12 in terms of line broadening and CSP profiles, suggesting indeed that the N-terminal part of the β1- strand and the α-helix constitute the true interaction surface (**Supplementary Figure S2**). Taken together, comparison of ^15^N CXCL12 NMR titrations with HMGB1 fragments revealed that the acidic IDR of HMGB1 is directly involved in HMGB1●CXCL12 heterocomplex formation and that the CXCL12 dimerization surface works as hub for multivalent interactions with both the acidic IDR and the HMG tandem domain.

### The HMGB1●CXCL12 heterocomplex forms via fuzzy interactions

We next hypothesized that the inter-molecular interactions between the acidic IDR and CXCL12 might perturb HMGB1 intra-molecular interactions and conformational equilibria. Indeed, in reversed titrations, HMGB1 amide resonances of *spy* residues reported to interact with the acidic IDR (e.g. W48, T76, I78 on BoxA, and A93, S016, I158 on BoxB)^30^, moved towards their NMR frequency in the tailless construct upon addition of CXCL12. These displacements suggest a weakening of the intramolecular interaction between the acidic IDR and the HMG boxes (**Figure 2A**). The shifts towards HMGB1-TL resonances were relatively small, presumably because CXCL12 only partially competes with HMGB1 intra-molecular interactions. Addition of an equimolar amount of Ac-pep to ^15^N HMGB1 in complex with CXCL12 was sufficient to sequester CXCL12 and disrupt the heterocomplex, as indicated by the reappearance of ^15^N HMGB1 resonances, confirming the important contribution of the acidic IDR to heterocomplex formation (**Figure 2B**).

**Figure 2:**
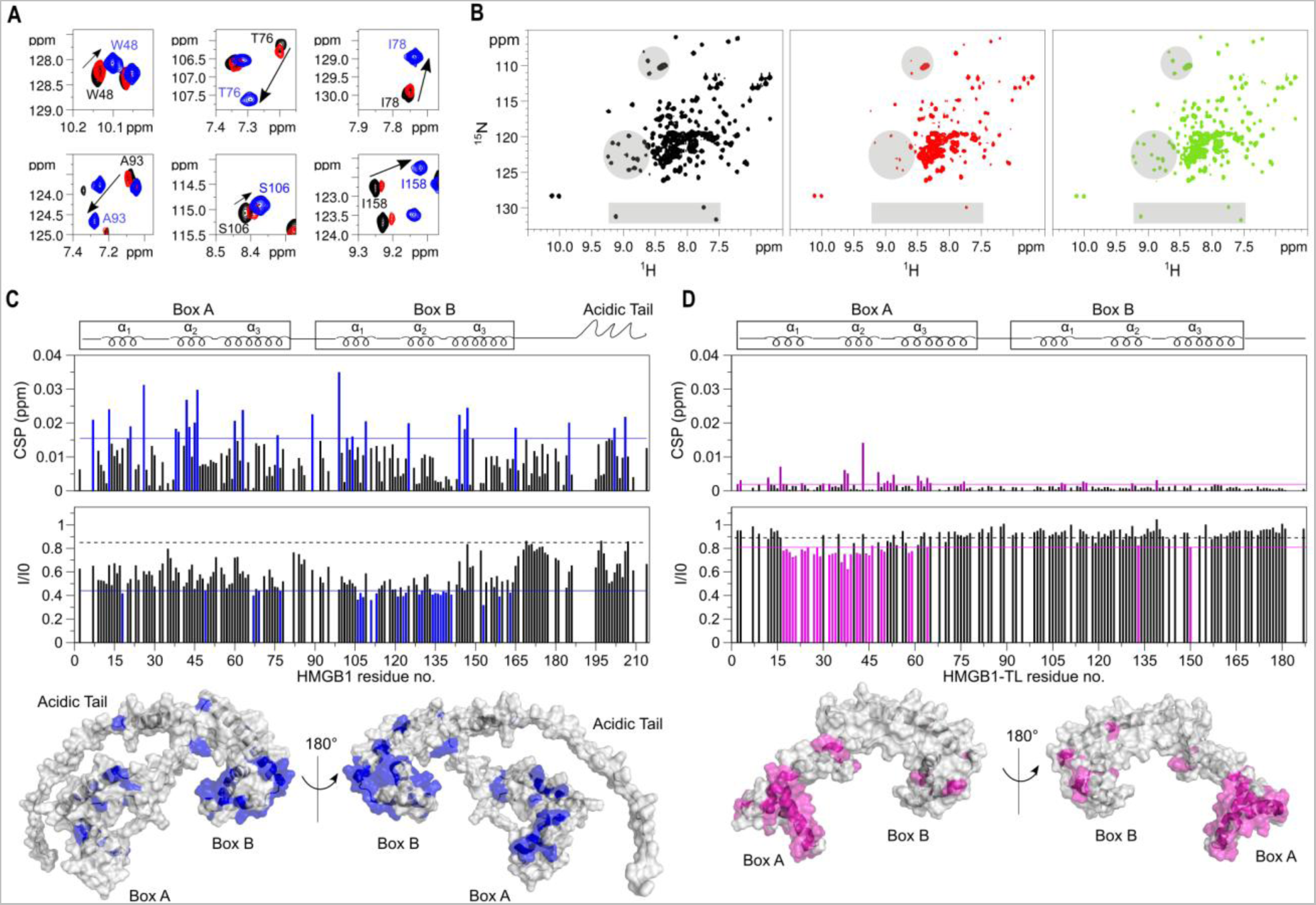
The HMGB1●CXCL12 heterocomplex forms via fuzzy interactions. **(A)** Superposition of selected regions of ^1^H-^15^N HSQC spectra of 0.1 mM ^15^N HMGB1 (corresponding to spy residues W48, T76, I78, A93, S106, I158) without (black) and with 0.2 mM CXCL12 (red) and 0.1 mM ^15^N HMGB1-TL (blue). CXCL12 partially competes with intramolecular HMGB1 interactions and specific amide resonances move (arrow) towards the chemical shift of the corresponding amide in the tailless construct. **(B)** ^1^H-^15^N HSQC spectra of HMGB1 (0.1 mM) without (black) and with 0.2 mM CXCL12 (red), and with subsequent addition of 0.1 mM Ac-pep (green). Grey shadowed regions highlight resonances disappearing and reappearing upon addition of CXCL12 and Ac-pep, respectively. **(C)** Bar graphs showing residue-specific CSPs (upper panel) and peak intensity ratios (I/I_0_) (lower panel) of ^15^N-labeled HMGB1 (0.1 mM) upon addition of CXCL12 (1:1). Residues with CSP > Avg+ σ_0_ (blue line) and (I/I_0_) < Avg - SD (blue line) are colored (blue) and mapped on HMGB1 (grey surface, alphafold2 model AF-P63158). **(D)** Bar graph showing residue-specific CSPs (upper panel) and intensities ratios (I/I_0_) (lower panel) of ^15^N-labeled HMGB1-TL (0.1 mM) upon addition of CXCL12 (1:1). Residues with CSP > Avg + σ_0_ (magenta line) and I/I_0_<Avg - SD (magenta line) are colored in magenta and mapped on HMGB1-TL (grey surface, pdb code: 2YRQ). In the bar-graphs α-helices are schematically represented on the top, missing residues are either prolines, or superimposed residues of the acidic IDR or absent because of exchange with the solvent, the dashed black line indicates the peak intensity decrease due to the titration dilution effect.

Overall, CSPs and intensity variations in ^15^N HMGB1/CXCL12 NMR titrations are due to both intra- and inter-molecular interactions, hence mapping of spectral perturbations on the HMGB1 structure reflects both phenomena, that are difficult to be separated (**Figure 2C**). Thus, to remove the “confounding” effect of the acidic IDR, we performed NMR titrations of ^15^N HMGB1-TL with CXCL12. Notably, the CSPs and peak intensity reduction profiles were different and substantially smaller than the ones observed in the full-length protein. Nonetheless, the removal of the acidic IDR brought out the presence of an interaction surface formed by the first two helices of BoxA, with no major involvement of BoxB (**Figure 2D**).

Taken together, NMR titrations indicate that HMGB1●CXCL12 is a *fuzzy* dynamic heterocomplex, characterized by multivalent inter- and intra-molecular equilibria involving CXCL12, the HMG tandem domain and the acidic IDR as major players within this intricate network of interactions.

### The acidic IDR of HMGB1 interacts with CXCL12 via long-range electrostatic interactions

We next adopted the same dissection approach to investigate the thermodynamics of CXCL12 interaction with HMGB1, using a combination of isothermal titration calorimetry (ITC), microscale thermophoresis (MST), and fluorescence measurements. ITC injection of CXCL12 into Ac-pep solution generated spikes with a biphasic profile, indicative of different binding events, and large maximal exothermic heat changes (∼-0.6 µcal/s). Global fitting of the buffer-subtracted binding isotherm yielded an apparent Kd_1_ of 0.6 ± 0.1 µM and Kd_2_ of 0.1 ± 0.1 µM, in agreement with the low micromolar affinity estimated by NMR line-shape analysis (**Figure 3A, Table 1**). Also the interaction between HMGB1 and CXCL12 appeared biphasic and exothermic (**Figure 3A**), though with one order of magnitude reduced amplitude (∼-0.06 µcal/). The fitting of the curve yielded an apparent Kd_1_ = 1.2 ± 0.4 µM and a second one, whose nanomolar value should be taken with caution because of the large error in the global fitting (**Table 2**). Importantly, the low micromolar affinities measured by ITC for Ac-Pep and HMGB1 were in good agreement with the ones deriving from fluorescence and MST experiments, respectively (**Figure 3B, Table 1**). While the heat of reaction between CXCL12 and HMGB1-TL was not sufficient to derive any binding parameters (endothermic spikes at baseline level, ∼0.05 µcal/s) (**Figure 3A**), a change of the thermophoretic diffusion properties of fluorescently labeled CXCL12 was detectable in the presence of HMGB1-TL. Notably, the derived affinity (Kd=12.5 ± 5.5 µM) was almost one order of magnitude weaker than the one measured with full-length HMGB1 (Kd=1.7 ± 0.2 µM), thus confirming the important role of the acidic IDR in heterocomplex formation (**Table 1**).

**Figure 3:**
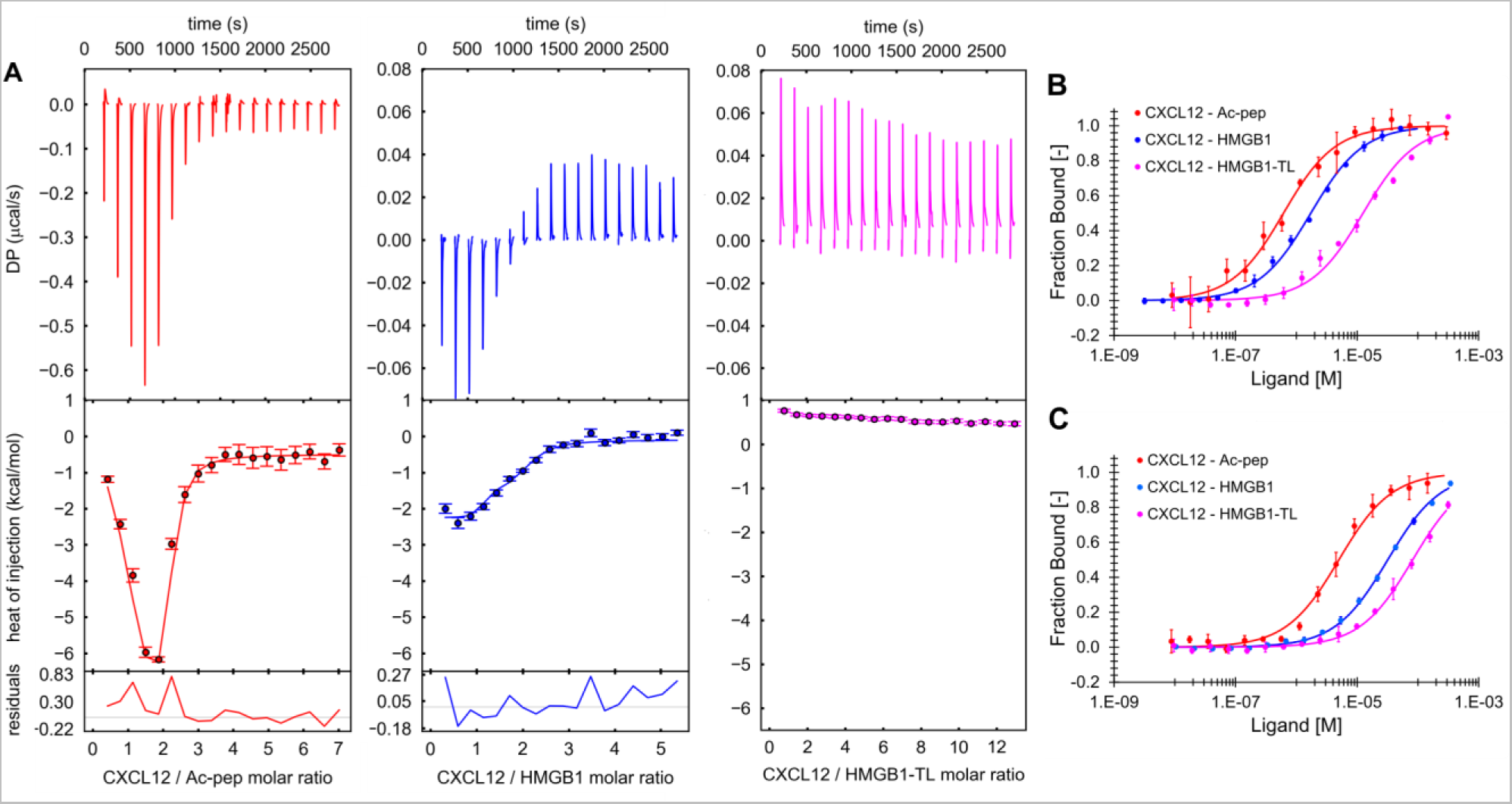
The acidic IDR of HMGB1 interacts with CXCL12 via long-range electrostatic interactions. **(A)** ITC measurements of CXCL12 titrated into Ac-pep (left, red), HMGB1 (middle, blue) and HMGB1-TL (right, magenta). The upper, middle and lower panels show respectively, the ITC sequential heat pulses for binding, the integrated data corrected for heat of dilution and fit to a two-site-binding model with a nonlinear least-squares method (line), and the residuals. Error bars indicate the error on the global fittings. One representative curve (n=3) for each titration is shown. Normalized variation of fluorescence of 5,6-FAM-labelled Ac-pep upon addition of CXCL12 and normalized variation of MST signal of labeled CXCL12 in the presence of HMGB1 (blue) and of HMGB1-TL (magenta), with 20 mM NaCl **(B)** and with 150 mM NaCl **(C).** In **(B)** and **(C)** n = 3; data represent Avg ± SD.

**Table 1:** Thermodynamic parameters of the interactions between CXCL12 and Ac-pep, HMGB1 and HMGB1-TL.

| **ITC** |  |  |  |
| --- | --- | --- | --- |
| **Param.<sup>1</sup>** | **Ac-pep** | **HMGB1** | **HMGB1-TL** |
| $Kd_1$ [$\mu M$] | $0.6 \pm 0.1$ | $1.2 \pm 0.4$ | *n.d.* |
| $Kd_2$ [$\mu M$] | $0.1 \pm 0.1$ | $7.8 \times 10^{-3} \pm 1.5 \times 10^{-2}$ | *n.d.* |
| $\Delta H_1$ [kcal/mol] | $-10.4 \pm 0.8$ | $-1.4 \pm 0.1$ | *n.d.* |
| $\Delta H_2$ [kcal/mol] | $1.6 \pm 0.7$ | $-2.1 \pm 0.2$ | *n.d.* |

| **MST** |  |  |  |
| --- | --- | --- | --- |
| **Param.** | **Ac-pep<sup>2</sup>** | **HMGB1<sup>3</sup>** | **HMGB1-TL<sup>4</sup>** |
| $Kd_{20mM, NaCl}$ [$\mu M$] | $0.6 \pm 0.1$ | $1.7 \pm 0.2$ | $12.5 \pm 5.5$ |
| $Kd_{150mM, NaCl}$ [$\mu M$] | $5.0 \pm 0.6$ | $31.8 \pm 1.4$ | $81.4 \pm 6.8$ |

<sup>1</sup> CXCL12 is the titrant, n=3, error estimates from covariance matrix<sup>2</sup> CXCL12 is the titrant, n=3, values are the average ± SD<sup>3</sup> HMGB1 is the titrant, n=3, values are the average ± SD<sup>4</sup> HMGB1-TL is the titrant, n=3, values are the average ± SD
n.d. not detected

**Table 2:** Calculated values from (A) AUC and (B) SAXS analysis of CXCL12, HMGB1 and HMGB1●CXCL12, pH 7.5, 20 mM TrisHCl, 50 mM NaCl.

| *C(s) model analysis of free components* |  |  |  | *MS-SV model analysis HMGB1: CXCL12 1:7.5* |  |  |
| --- | --- | --- | --- | --- | --- | --- |
|  | *Param.* | **CXCL12** | **HMGB1** | *Param.* | **CXCL12<sub>mix</sub>** | **HMGB1<sub>mix</sub>** |
| **(A) AUC-SV** | $sw^1$ | 1.1 S | 2.3 S | $sw^5_{(20,w)}$ | 2.8 S | 3.1 S |
| | $f/f_0^2$ | 1.3<br>(globular) | 1.4<br>(elongated) | $conc^6_{tot}$ | 43 μM | 7.7 μM |
| | $MW (kDa)^3$ | 8.4 | 27.1 | $conc^7_{complex}$ | 8.5 μM | 7.7 μM |
| | $RMSD^4$ | 0.0052 | 0.0051 | $RMSD^8_{280nm}$<br>$RMSD^9_{250nm}$<br>$D_{norm}^{10}$ | 0.0072,<br>0.0050<br>0.08 | |
| **(B) SAXS** | $R_g (nm)^{11}$ | 1.5 ± 0.01 | 2.6 ± 0.02 | $R_{g, complex} (nm)^{11}$ | 2.9 ± 0.02 | |
| | $D_{max} (nm)^{12}$ | 4.8 | 8.5 | $D_{max, complex}^{12}(nm)$ | 10.5 | |
| | $MW (kDa)^{13}$ | 9.7 | 26.4 | $MW_{complex} (kDa)^{13}$ | 35.8 | |

<sup>1</sup>sedimentation coefficient<sup>2</sup>frictional ratio, f is the frictional parameter of the particle and $f_0$ is the frictional value of a smooth sphere<sup>3</sup>estimated molecular weight<sup>4</sup>RMSD, root mean square deviation of the fitting<sup>5</sup>sedimentation coefficient normalized to standard solution conditions of water at 20° C expressed in Svedberg (S)<sup>6</sup>concentration of the components; $conc_{tot}$ <sup>7</sup>calculated concentration of the component in the complex; $conc_{complex}$ <sup>8</sup>root mean square deviation of the fitting at 280 nm<sup>9</sup>root mean square deviation of the fitting at 250 nm<sup>10</sup> $D_{norm}$ determinant of the extinction coefficient matrix.<sup>11</sup>radius of gyration<sup>12</sup>Maximum distance of the pairwise distance distribution plots<sup>13</sup>mass estimate based on volume

We reasoned that the interaction could be dominated by long range electrostatic interactions between the acidic IDR and the basic surface of CXCL12 (**Supplementary S3A, B**). Indeed, the heat of reaction associated to CXCL12 interaction with Ac-pep or HMGB1 at higher ionic strength (150 mM NaCl) did not yield any spikes above baseline (**Supplementary Figure S3C, D**). Also in fluorescence and MST experiments binding affinities were reduced by one order of magnitude, confirming the prominent contributions of long-range electrostatic interactions (**Figure 3C, Table 1**).

Collectively, ITC, MST, and fluorescence measurements, in accordance with NMR titrations, support a scenario in which the HMG tandem domain and the acidic IDR both contribute to heterocomplex formation, with the acidic IDR working as an “antenna” for recruiting CXCL12 via long-range electrostatic interactions.

### HMGB1 and CXCL12 form an equimolar heterocomplex

Previously, we and others postulated that HMGB1 and CXCL12 interact with a 1:2 ratio ^26, 27^. To verify this hypothesis, we rigorously assessed complex stoichiometry by analytical ultracentrifugation (AUC). Initially, we compared sedimentation velocity AUC (SV-AUC) experiments of the individual and combined components at different stoichiometric ratios. Sedimentation coefficients (c(S)) of free CXCL12 (1.1 S) and HMGB1 (2.3 S) were well in agreement with their associated apparent molecular weights (8.7 and 27KDa, respectively) (**Figure 4A, B; Table 2A**), and the corresponding frictional ratios (f/f_0_) of 1.3 and 1.4 were in line with the globular and elongated shape of CXCL12 and HMGB1, respectively (**Table 2**). At variance to stable complexes, characterized by distinct SV-AUC curves for the bound and individual components, the HMGB1●CXCL12 heterocomplex presented only two separable sedimentation distributions: one relatively sharp peak at lower c(S), corresponding to free CXCL12, and one at higher c(S) values, deriving from the sedimentation of free and bound HMGB1. The latter, both at 50 (**Figure 4C, Table 2A)** and 150 mM NaCl (**Supplementary Figure S4A-C, Supplementary Table S1**) was relatively broad and shifted towards higher c(S) values (and apparent MWs) with increasing CXCL12 concentrations. This behavior is typically observed in highly dynamic complexes, where the reaction boundaries between bound and unbound species cannot be resolved within the signal-to-noise of the experiment^31^. Importantly, analysis of SV-AUC curves of HMGB1-TL with increasing concentrations of CXCL12 showed only two peaks corresponding to the free components, and no sedimentation peaks displacement or broadening were observed. Conceivably, without the acidic IDR the association between the two components is too rapid to be detected in the sedimentation time scale (**Supplementary Figure S4D-F).**

**Figure 4:**
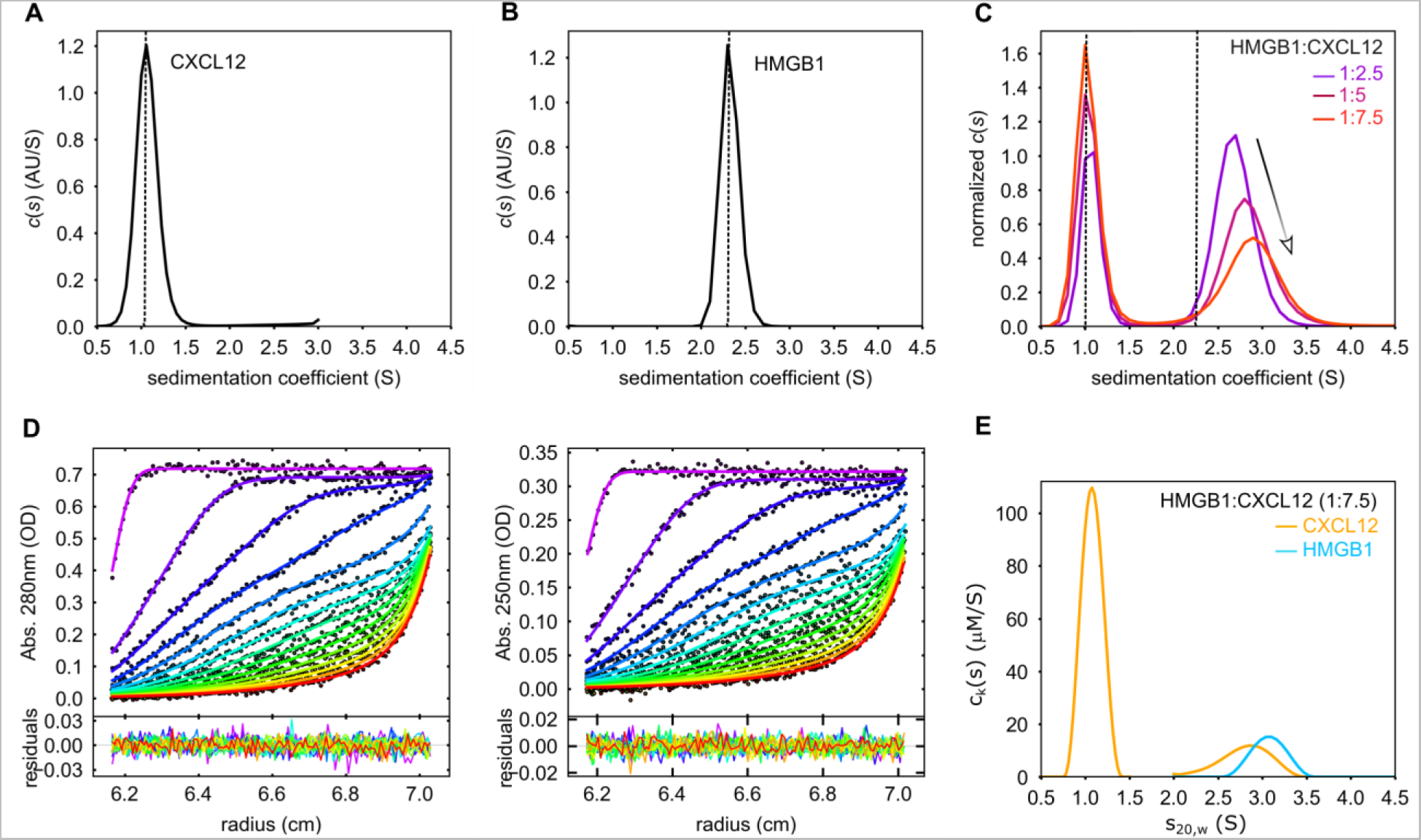
HMGB1 and CXCL12 form a transient 1:1 heterocomplex. Sedimentation velocity analytical ultracentrifugation (SV-AUC) experiments of free (**A**) CXCL12 (7.7 µM) and (**B**) HMGB1(7.7 µM), scanned by absorbance at 280nm. (**C**) SV-AUC analysis of the interaction between HMGB1 (7.8 µM) and increasing concentrations of CXCL12 (colors). The dotted lines indicate the sedimentation coefficients of the free components. (**D**) Global multi-signal sedimentation velocity (MS-SV) analysis to determine the stoichiometry of HMGB1:CXCL12 complex, with 7.7 µM HMGB1 and 43 µM CXCL12. The raw sedimentation signals of HMGB1:CXCL12 mixture acquired at different time points with absorbance at 280 nm (left), and absorbance at 250 nm (right) with the corresponding signal profiles as a function of radius in centimeters. The time-points of the boundaries are indicated in rainbow colors, progressing from purple (early scans) to red (late scans). Only every 3rd scan used in the analysis are shown. Residuals of the fit are shown at the bottom. (**E)** Decomposition into the component sedimentation coefficient distributions, ck(s), for CXCL12 (yellow line) and HMGB1 (cyan line).

Next, to determine the ratio of HMGB1 and CXCL12 in the complex distribution, we performed multi-signal SV-AUC experiments (MS-SV-AUC) and exploited the different extinction coefficients of HMGB1 and CXCL12 at minimal (250 nm) and maximal (280 nm) wavelengths to distinguish between the two complex components^32, 33^. Analysis of MS-SV-AUC suggested that HMGB1 predominantly forms with CXLC12 (or CXCL12-LM) a 1:1 heterocomplex in solution, as indicated by the equimolar concentrations for both CXCL12 (CXCL12-LM) and HMGB1 obtained from the peak area at ∼3S (**Figure 4D, E, Table 2A, Supplementary Figure S4G-K, Supplementary Table S2**).

Based on these experiments we conclude that HMGB1 and CXCL12, in contrast to previous assumptions, form an equimolar heterocomplex, with the acidic IDR playing a fundamental role in complex assembly.

### Small Angle X-ray scattering (SAXS) analysis supports the dynamic nature of the HMGB1●CXCL12 heterocomplex

We then utilized small angle X-ray scattering (SAXS) to obtain low-resolution structural information on the mass and shape of the heterocomplex. Primary analysis of the SAXS scattering curves of CXCL12 and HMGB1 yielded radii of gyration (R_g_) of 1.55 nm and 2.60 nm, respectively, as well as pairwise distance distribution plot (P(r)) derived maximum distance (D_max_) values of 4.8 nm and 8.6 nm. These values indicate that both proteins are monomeric and monodisperse in solution. (**Figure 5A, Table 2B, Supplementary Figure S5, Supplementary Table S3**). The values obtained for HMGB1 are in accordance with a previous report^23^. The normalized Kratky representation of free CXCL12 and HMGB1 displayed upward trends at higher qR_g_ values well in agreement with the presence of flexible regions within a folded core (**Figure 5B**). Analysis of the heterocomplex was challenging due to its transient and dynamic nature, which prevented its isolation through size exclusion chromatography and direct coupling to SAXS. Instead, we used a “batch-mode” strategy^34^ where HMGB1 incubated with increasing CXCL12 concentrations was immediately analyzed by SAXS. The molecular dimension of a 1:1 complex with an estimated molecular weight of 35 kDa was obtained in the presence of two CXCL12 equivalents, suggesting a dynamic exchange between bound and unbound CXCL12 (**Table 2B**). At higher CXCL12 concentrations we could not estimate reliably the molecular weight due to unbound CXCL12 (**Supplementary Figure S5A**). Thus, we focused our SAXS analysis on the sample comprising two CXCL12 equivalents. We observed an overall increase of the derived parameters with respect to free HMGB1, compatible with complex formation (R_g_ of 2.90 nm and D_max_ of 10.2 nm) and a mass estimation corresponding to HMGB1●CXCL12 heterocomplex. The normalized Kratky plot confirmed the dynamic and flexible nature of the complex, containing both folded domains and unstructured segments (**Figure 5B**). Next, to model more quantitatively the fuzzy nature of HMGB1●CXCL12 heterocomplex, we combined NMR-guided rigid-body docking (SASREF^35^ and FoXSdock^36^) to enhanced optimization methods (EOM)^37^ (details in materials and methods). Herewith, we identified two main conformational ensembles fitting well the experimental data (with χ^2^ <1, **Supplementary Table S3**). In the first ensemble, CXCL12 binds to BoxA of HMGB1, adopting alternatively open or more collapsed conformations (**Figure 5C**). In the second one, the acidic IDR of HMGB1 wraps around CXCL12, while the two HMG boxes adopt different reciprocal orientations (**Figure 5D**).

**Figure 5:**
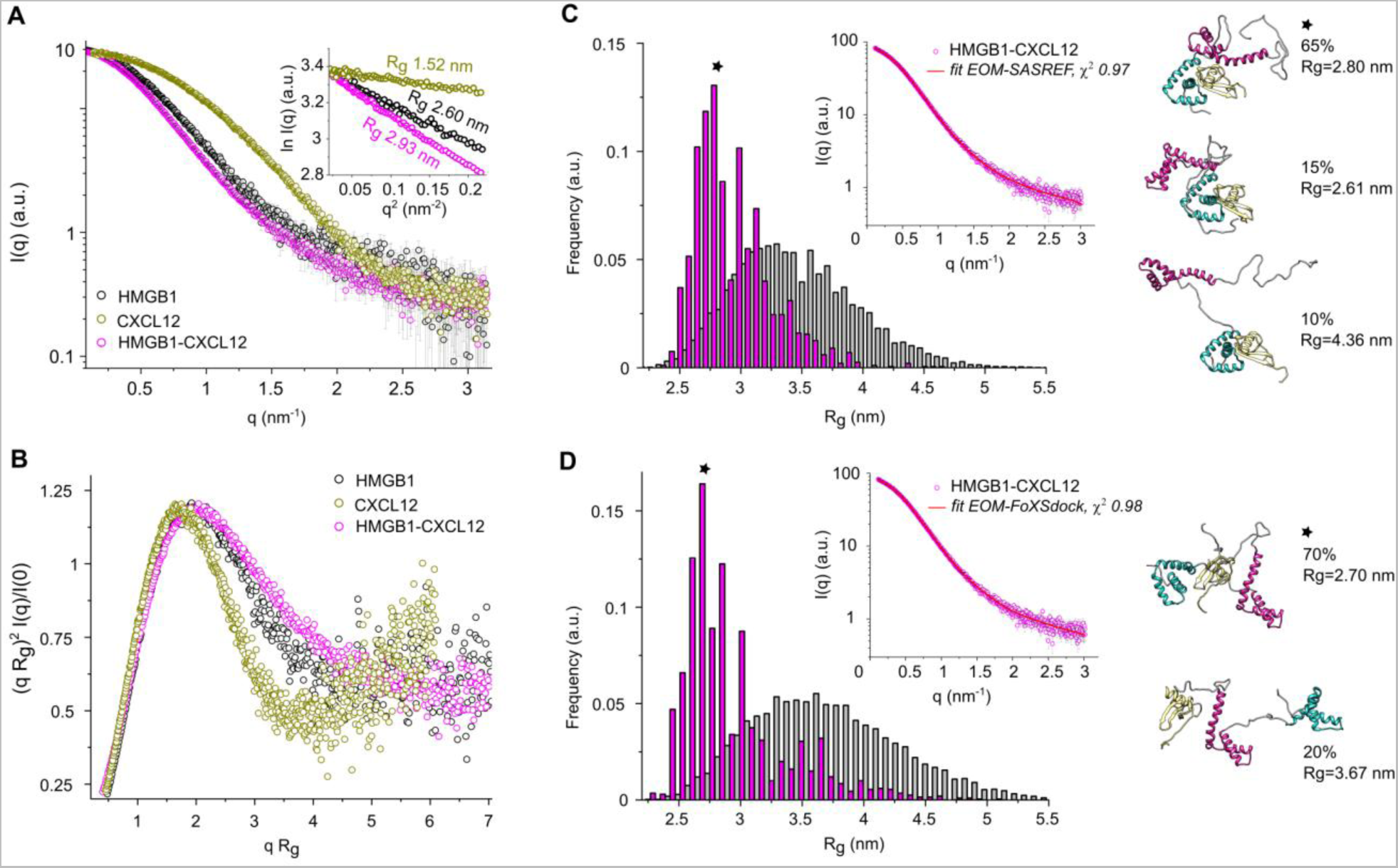
SAXS studies of free HMGB1, CXCL12 and of HMGB1●CXCL12. **(A)** I(*q*) *versus q* experimental SAXS profiles for CXCL12 (gold), HMGB1 (black) and HMGB1●CXCL12 complex (magenta). Error bars represent an estimate of the experimental error σ on the intensity recorded for each value of *q* assigned by data reduction software. In the inset, the Guinier regions used to estimate the radii of gyration (R_g_, nm). **(B)** Dimensionless Kratky plots for the data presented in **(A)**. Analysis of SAXS data by EOM on HMGB1●CXCL12 models obtained using **(C)** SASREF and **(D)** FoXSdock with distributions of the selected ensemble conformers (magenta bars) and the initial pools of structures (grey bars) as a function of R_g_ in nm. In the insets, I(*q*) *versus q* (magenta squares) with the EOM fitting (red lines with the corresponding χ^2^ values) for the HMGB1●CXCL12 complex. Representative structure of the most populated EOM ensembles are shown in cartoon, with BoxA, BoxB, IDR and CXCL12 coloured in cyan, magenta, grey and gold, respectively. For each ensemble, the frequency-weighted size average (the asterisks indicate the most populated fractions) and R_g_ values are indicated.

Overall, SAXS analysis supports the notion that HMGB1 and CXCL12 form an equimolar *fuzzy* complex.

### The acidic IDR modulates the interaction of the HMGB1●CXCL12 heterocomplex with the CXCR4 receptor

Having shown that the acidic IDR of HMGB1 plays a major role in the interaction with CXCL12, we tested whether it also played a role in the binding of the heterocomplex to its CXCR4 receptor. For these experiments, we chose mouse AB1 cells, a cellular model of malignant mesothelioma^38^, as they express high levels of CXCR4^39^.

We first verified the existence of the HMGB1●CXCL12 heterocomplex in association with CXCR4 on the cell membrane. Proximity ligation assays (PLA) between HMGB1 and CXCL12 clearly identified the HMGB1●CXCL12 heterocomplex on the surface of AB1 cells, and the PLA signal was competed by increasing concentrations of AMD3100, a specific CXCR4 antagonist **(Figure 6A)**, confirming that the heterocomplex binds CXCR4. Likewise, treatment of cells with increasing amounts of Ac-pep significantly decreased the amount of detectable heterocomplex, in line with the observation that Ac-pep can bind CXCL12 and disrupts the HMGB1●CXCL12 heterocomplex.

**Figure 6:**
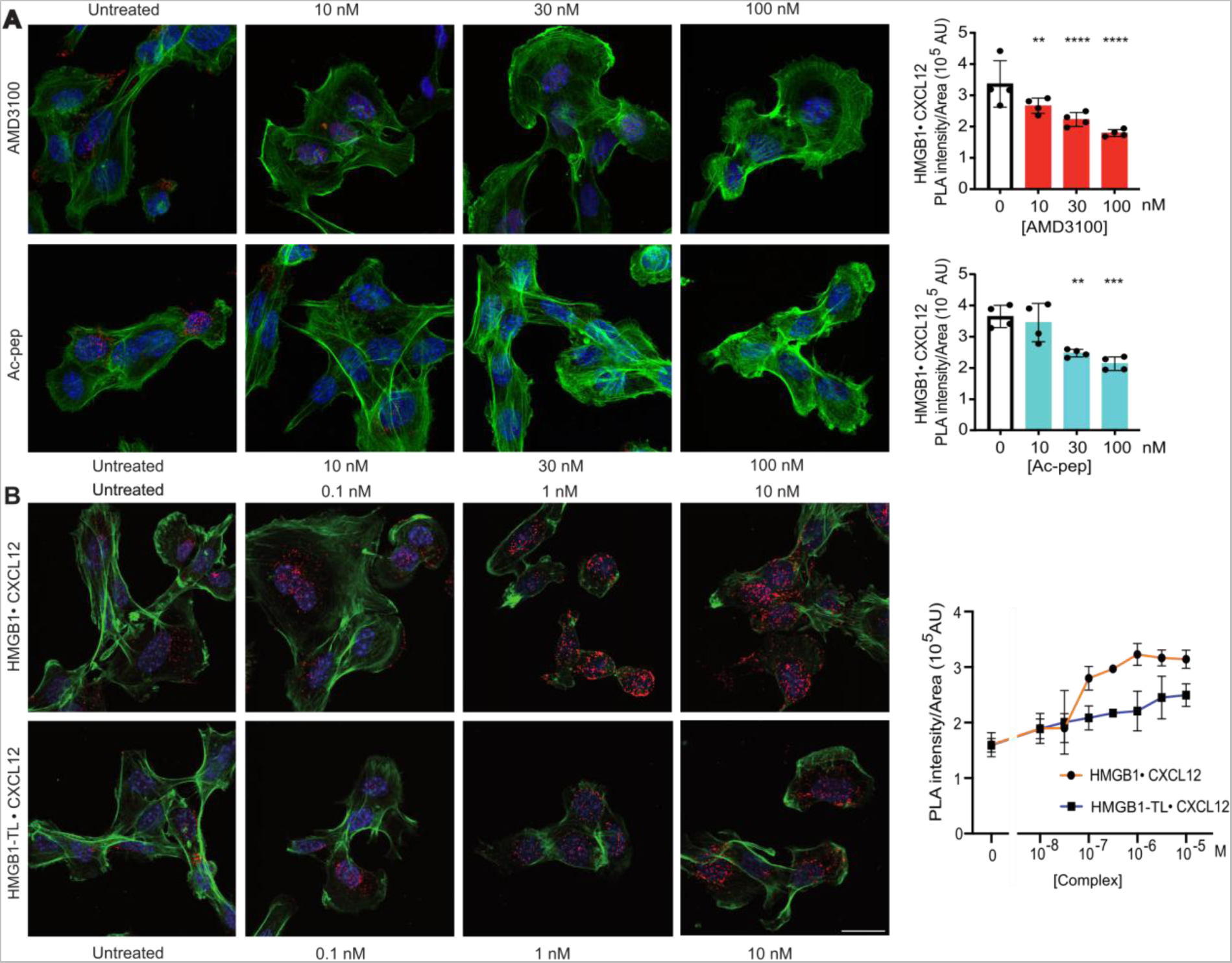
The acidic IDR modulates HMGB1●CXCL12 binding to CXCR4 on AB1 cells. **(A)** Representative confocal microscopy images of Proximity Ligation Assays (PLAs) performed on the HMGB1•CXCL12 complex on the surface of AB1 malignant mesothelioma cells. Cells were either untreated or treated with AMD3100 or Ac-pep for 1 hour at 37°C/5% CO_2_ at three different concentrations (10, 30, or 100 nM). Nuclei are in blue (Hoechst 33342), phalloidin is in green, and the HMGB1•CXCL12 PLA signal is red. Scale bar; 20 μM. PLA signal was quantified as described in the Methods section. One-way ANOVA was performed comparing Ac-pep treated cells to untreated cells; \*\**P* <0.01; \*\*\**P* <0.001, \*\*\*\**P* <0.0001. **(B)** PLA signal quantification on the surface of ligand-stripped AB1 cells exposed at 4°C to increasing concentrations of either HMGB1•CXCL12 or HMGB1-TL•CXCL12 equimolar heterocomplexes. Colors and PLA as in panel **(A)**. Mean ± SD are indicated; n=4 to 6 per concentration. The difference between HMGB1•CXCL12 and HMGB1-TL•CXCL12 heterocomplexes is statistically significant (*P* <0.0001; two-way ANOVA).

We then tested the binding to cell surface CXCR4 of heterocomplexes made with full-length or tailless HMGB1 (HMGB1-TL). Cells were washed at acidic pH to remove ligands bound to receptors, and then exposed to preformed mixtures of CXCL12 and full-length or HMGB1-TL; cells were kept at 4°C to prevent receptor internalization. Both full-length and tailless HMGB1 formed complexes that could be detected by PLA on the cell surface; however, HMGB1-TL formed fewer complexes and did not reach saturation (**Figure 6B**).

Together, these results show that the HMGB1 acidic IDR facilitates the binding of the HMGB1●CXCL12 heterocomplex to its CXCR4 receptor, and that the Ac-pep competes with the acidic IDR for heterocomplex formation and binding to CXCR4.

## Discussion

Here, we shed new light on the molecular mechanisms at the basis of HMGB1●CXCL12 heterocomplex formation, and suggest a major role of fuzzy interactions mediated by the acidic IDR of HMGB1. Importantly, we also show that the stoichiometry of the HMGB1●CXCL12 heterocomplex is 1:1.

Until now, structural studies aimed at getting mechanistic insights into HMGB1●CXCL12 have focused on the interaction of CXCL12 with the single isolated HMG boxes^15, 26^. The question arose whether such a simplified approach, in which the HMG boxes are separated and the IDRs (i.e. the acidic C-terminal tail and the linker connecting the HMG boxes) are neglected, can faithfully recapitulate the interaction of CXCL12 with the full-length protein. IDRs within their host proteins are often the major players in the recognition of their partners^40–42^; this statement also applies to HMGB1, where the acidic C-terminal IDR modulates interactions with nucleic acids^43^ and proteins, such as histones^44, 45^ and p53^46^. Importantly, replacement of the acidic IDR with an arginine-rich basic tail has been recently shown to cause a complex human malformation syndrome, which results from HMGB1 aberrant phase separation in the nucleolus and nucleolar dysfunction^47^. In accordance with its functional relevance, here we show that the acidic IDR, whether isolated or in the context of the full-length protein, binds to the CXCL12 homodimerization surface. Noteworthy, binding of full-length HMGB1 to CXCL12 is significantly weaker than to the isolated acidic IDR, indicating competition of the intra-molecular HMGB1 boxes with CXCL12 for the acidic IDR. Both ITC and MST experiments suggest long range electrostatic interactions between the acidic IDR and the basic surface of CXCL12 as major drivers for binding, as increased ionic strength or deletion of the acidic IDR reduced binding to CXCL12. Long-range electrostatic interactions, though fundamental, are not the unique driving force for complex formation, as both NMR and MST indicate that tailless HMGB1 binds with micromolar affinity to the dimerization surface of CXCL12, albeit less with respect to the full-length protein. Thus, structured and unstructured H0MGB1 regions potentially recognize the same CXCL12 surface, which behaves as a structural hub for multivalent interactions. Notably, while the isolated HMG-boxes were previously suggested to interact similarly with CXCL12^15^, a preference for BoxA is apparent in the context of the tandem HMG-boxes, with the interaction surface mainly involving the two short helices of the domain. Of note, the preferential targeting of BoxA is in line with the ability of CXCL12 to recognize the reduced forms of Cys-22 and Cys-44^15, 48^. Preferential binding to BoxA is also in agreement with the equimolar stoichiometry of the heterocomplex suggested by both AUC and SAXS experiments.

Collectively, our data reveal that the HMGB1●CXCL12 heterocomplex behaves as a typical *fuzzy complex*^49^, whose formation relies on an intricate network of inter- and intra- molecular interactions of comparable affinities. As commonly observed in fuzzy binding, at least one of the elements –in this case the acidic IDR– is dynamic and fundamental for the interaction. In addition, the intrinsic independent rotation of the two HMG-boxes provides an additional dynamic level to the system. Thus, the heterocomplex cannot be described by a unique structure, but is best represented by a heterogeneous ensemble of structures reflecting the different ongoing equilibria. Accordingly, the SAXS data of the heterocomplex are best fit by different plausible docking models obtained by EOM, where CXCL12 binds HMGB1 in a promiscuous manner, alternatively associating to the manifold conformations of the acidic IDR and the different BoxA orientations.

Based on our data we propose a model (**Figure 7**) in which the acidic IDR works as a wrapping antenna that recruits cellular CXCL12 through long-range electrostatic interactions. Being intrinsically disordered, the acidic IDR does not present a single binding site to CXCL12 but rather resembles a diffuse ‘‘binding cloud’’, in which multiple nearly-identical binding sites are dynamically distributed ^50, 51^, preserving a significant flexibility even in bound states. CXCL12 then can interact with HMG-boxes, in particular with BoxA in its reduced form, partially outcompeting HMGB1’s intramolecular contacts. Remarkably, the binding mode of HMGB1 to CXCL12 radically differs from the usual beta-beta or alpha-beta interactions observed in chemokine heterocomplexes^5^.

**Figure 7:**
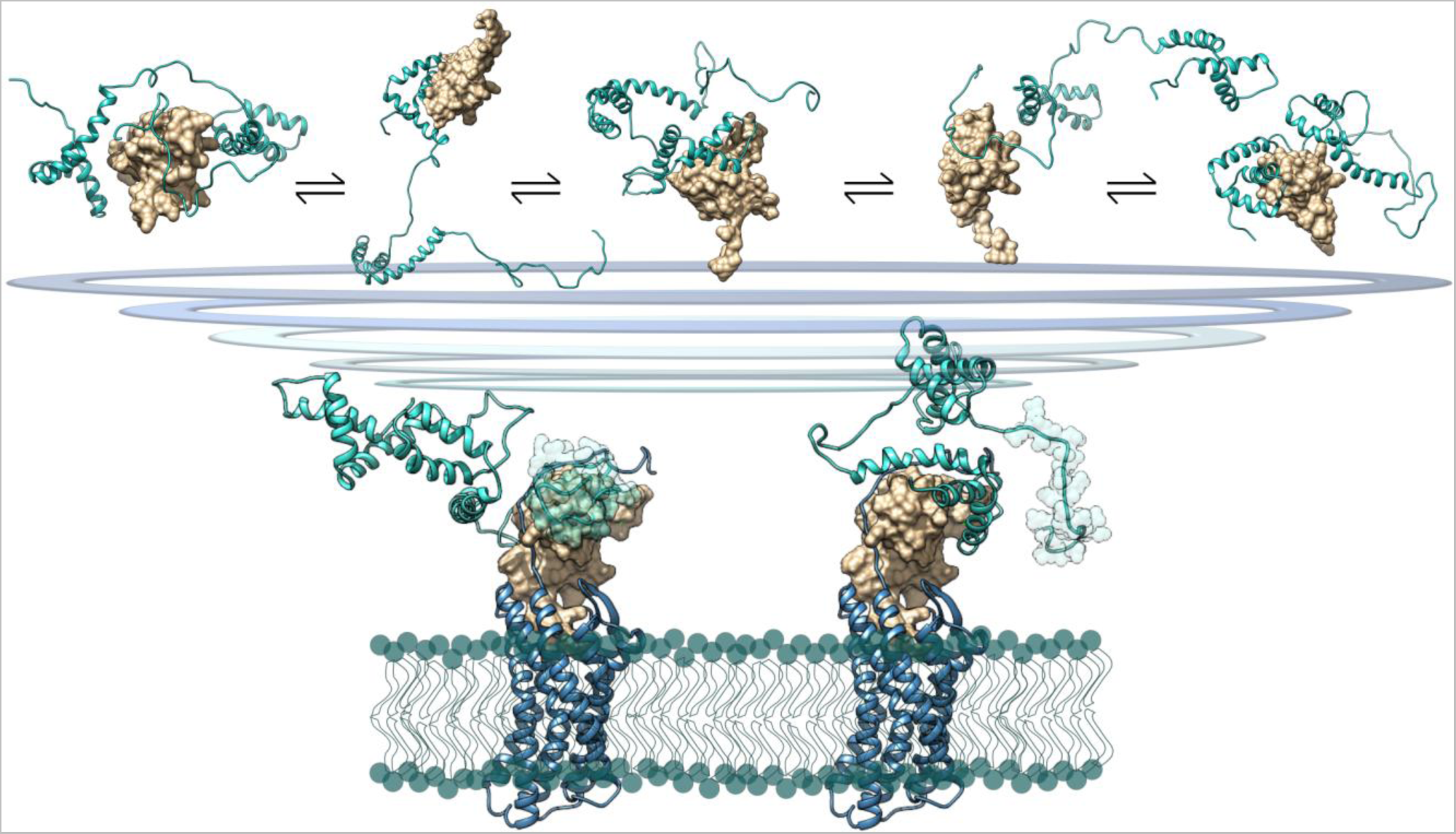
Model of HMGB1●CXCL12 fuzzy complex. *Top*: Model of the *fuzzy* interactions between HMGB1 (cyan cartoon) and CXCL12 (gold surface) *Bottom*: explicative representations of possible different CXCL12●HMGB1 conformations bound to CXCR4 (blue cartoon). Two SAXS-EOM HMGB1●CXCL12 models have been superimposed on the theoretical model of CXCL12 in complex with CXCR4^80^, with CXCL12 represented in gold surface and HMGB1 in cyan, the acidic IDR is highlighted with spheres; the lipid bilayer is represented with green spheres and lines.

CXCL12 bound to HMGB1 can then accommodate inside the cradle formed by the transmembrane helices of CXCR4, yielding a three-component complex (**Figure 7**), whose existence on the cell surface is supported by PLA experiments. These results are in line with the observation that the HMGB1●CXCL12 heterocomplex acts differentially from CXCL12 alone on the reorganization of the actin cytoskeleton and on β−arrestin recruitment^21^. Both actions depend on CXCR4. So far, however, our data do not specify whether a specific subset of the conformations of the HMGB1●CXCL12 heterocomplex preferentially binds to CXCR4. In our cellular context, the acidic IDR facilitates the binding of the HMGB1●CXCL12 heterocomplex to CXCR4, and Ac-Pep does compete with it, confirming the results obtained by NMR (**Figure 2**), and implying that the acidic IDR plays a major role in the formation of the CXCR4●HMGB1●CXCL12 ternary complex as well. The fuzziness of the interactions in the HMGB1●CXCL12 heterocomplex forces us to reconsider the structure of HMGB1, which itself can be considered fuzzy: in solution HMGB1 populates an ensemble of different micro-states in which the D/E repeats are associated through electrostatic transient interactions with different segments of HMGB1, that in turn are partially screened and only transiently exposed to natural interactors^52^. Plausibly, the fuzzy conformation of HMGB1 allows interactions with multiple partners^53^, all of which have micromolar-range apparent affinities. Indeed, HMGB1 often works as a chaperone, by binding one interactor and facilitating its further interaction with another molecule^54–56^. As such, the mechanism and the conformational heterogeneity through which HMGB1 binds to CXCL12 is in part reminiscent of other chaperones, like small heat shock proteins, that exploit their large IDR to transiently interact with their clients^57^.

In brief, we show here that HMGB1 and CXCL12 form a bimolecular heterocomplex, which is fuzzy and highly dynamical, and can go on to form a ternary complex with the CXCR4 receptor. These conclusions are in line with the concepts of HMGB1 working as a molecular chaperone and of IDPs/IDRs being typically involved in signaling, regulation, recognition, and control of various cellular pathways ^58^. We anticipate that interfering with the fuzzy interactions within the HMGB1•CXCL12 heterocomplex could represent a pharmacological strategy to inhibit its detrimental activity in inflammatory conditions ^59^.

## Methods

### Protein production and synthetic peptide

Recombinant labeled and unlabelled (^15^N/^13^C) HMGB1 (accession code P63158, residues 1–215) and HMGB1-TL (residues 1–187) were transformed in BL21 (DE3) pLysS and BL21 (DE3) strains of *Escherichia coli*, respectively using the expression vector pETM-11 vector (EMBL, Heidelberg, DE). The proteins were purified as described^60^. Recombinant labeled and unlabelled ^15^N/^13^C CXCL12 (accession code P48061, residues 1–68) was transformed into *Escherichia coli strain* BL21 (DE3) using the expression vector pET30a. The protein was purified as described in^61^. Unlabeled N-terminal-6His-tagged CXCL12 for MST measurements was provided by HMGBiotech (Milan, Italy).

For the production of CXCL12-LM site-directed mutagenesis was performed to introduce mutations L55C and I58C in CXCL12 pET30a expression vector by using standard overlap extension methods. The DNA constructs were sequenced by Eurofins (Milan, Italy). CXCL12-LM was expressed and purified as described^61^. Protein concentrations were determined considering molar extinction coefficients at 280 nm of 21430 and 8730 M^−1^ cm^−1^ for HMGB1 (and HMGB1-TL) and CXCL12, respectively.

Ac-pep (corresponding to HMGB1 acidic tail, 30 aa, residues 186-214) was purchased from Caslo Lyngby, Denmark. Peptide purity (>98%) was confirmed by HPLC and mass spectrometry. Peptide concentration was estimated from its dry weight. For ITC measurements, to obtain a more accurate estimation of the concentration by UV a tyrosine (ε_274nm_= 1,405 M^−1^ cm^-1^) was added at the peptide N-terminus, for MST experiments 5,6 FAM was added at the peptide N-terminus.

### NMR spectroscopy

NMR experiments were performed at 298K on a Bruker Avance 600 MHz equipped with inverse triple-resonance cryoprobe and pulsed field gradients (Bruker, Karlsruhe, Germany). Typical samples concentration was 0.1–0.4 mM. Data were processed using NMRPipe^62^ or Topspin 3.2 (Bruker) and analyzed with CCPNmr Analysis 2.4^63^. The ^1^H, ^13^C, ^15^N chemical shifts of CXCL12 in the presence of Ac-pep and of CXCL12-LM were obtained from three-dimensional HNCA, CBCA(CO)NH, CBCANH, HNCO experiments.

#### NMR Titrations

Before NMR titrations the samples (titrant and titrated solution) were dialyzed against the same buffer, 20 mM NaH_2_PO_4_/Na_2_HPO_4_ pH 6.3, 20 mM NaCl, supplied with 0.15 mM 4,4-dimethyl-4-silapentane-1-sulfonic acid (DSS) and D2O (10% v/v). In the case of Ac-pep titrations the lyophilized peptide was dissolved directly in the NMR buffer. Titrations were carried out by adding to ^15^N labelled protein samples (typically 0.1 mM) small aliquots of concentrated (15 mM) peptide stock solutions or unlabelled protein (0.6-1 mM). For each titration point (0.5, 1, 1.5, 2 equivalents of ligand) a 2D water-flip-back ^15^N-edited HSQC spectrum was acquired with 2,048 (160) complex points, apodized by 90◦ shifted squared (sine) window functions and zero filled to 2048 (512) points for ^1^H (^15^N). Spectra assignment was made following individual cross-peaks through the titration series. For each residue the weighted average of the ^1^H and ^15^N chemical shift perturbation (CSP) was calculated as 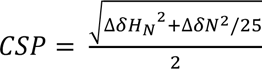 ^64^, where ΔδH_N_ and ΔδN are, respectively, the differences of ^1^H_N_ and ^15^N chemical shifts between free and bound protein. The corrected standard deviation (σ_0_) was calculated as described in^65^. Because of extensive line broadening due to ligand binding in the intermediate exchange regime on the NMR time scale, we also monitored changes in the intensity ratio (I/I_0_) of the ^1^H_N_-^15^N amide resonances, where I_0_ and I are the peak intensities in the free and bound protein, respectively.

#### Lineshape analysis

We performed 2D NMR lineshape analysis using the software TITAN^28^. Spectra were processed with NMRpipe^62^ with a script provided by TITAN. A series of regions of interest (ROI) containing isolated peaks with CSPs > Avg + SD were selected (V18, V23, K24, H17, A40, H25 and R12 for ^15^N-CXCL12 and V23, K24, H25 for ^15^N-CXCL12_LM) (**Supplementary Figure 1B,C**) and fitted by optimizing the chemical shifts and line widths for the free and the bound state. The program estimates *K*_d_, *k*off (we fixed 1:1 stoichiometry). Error estimates for the fit-parameters were obtained using the bootstrap resampling of residuals procedure implemented in TITAN^28^.

### Isothermal Titration Calorimetry (ITC)

Proteins and peptides were dialyzed in a Slide-A-lyser mini-dialysis unit with a 2,000 MWCO and Biodialyzer with 500 Da MWCO (Harvard Apparatus, US) against 20 mM TrisHCl at pH 7.5, 50 mM NaCl (or 150 mM NaCl when explicitly stated). ITC data were collected on a MicroCal PEAQ-ITC instrument (Malvern). The cell temperature was set to 37°C, the syringe stirring speed to 750 rpm, and reference power to 10 μcal/sec. HMGB1 (HMGB1-TL, Ac-pep) and CXCL12 were loaded into the cell and syringe at concentrations of ∼10 and ∼600 μM, respectively. The MicroCal PeakITC software (Malvern) was applied for initial data analysis. For global fitting, thermograms were integrated using NITPIC^66^ and SEDPHAT^67^. Data were fit with the two non-symmetric sites microscopic K model^68^ applying Simplex and Marquardt-Levenberg optimization algorithms, as implemented in SEDPHAT. Error estimates were based on the covariance matrix generated by the Marquardt-Levenberg algorithm.

### Microscale Thermophoresis (MST)

MST experiments were performed at 24°C on a NanoTemper® Monolith NT.115 instrument. Binding between Ac-pep and CXCL12 was monitored titrating CXCL12 (16- points) into 50 nM N-terminal-5,6-FAM-labelled Ac-pep (CASLO ApS, Denmark), using the blue filter, 20% LED power and medium MST power. Binding between HMGB1 or HMGB1-TL and CXCL12 was monitored titrating HMGB1 constructs (16-points) into 50 nM 6His- tagged CXCL12, non-covalently labelled with the NT-647 conjugated tris-NTA (RED-tris-NTA) fluorescence dye, using the red filter, 40% LED power and medium MST power. Before MST titrations the proteins (the ligand and the fluorescently labelled target) were dyalized against the same buffer, 20 mM NaH2PO4/Na2HPO4 pH 7.3, 0.05% TWEEN and 20 mM or 150 mM NaCl. In the case of 5,6-FAM-labelled Ac-pep, the lyophilized peptide was dissolved directly in the MST buffer and the pH adjusted to pH 7.3.

The 16 titration points of each experiment were made through serial dilution of the ligand stock into MST buffer and then addition of a constant amount of fluorescently labelled target (50 nM). Before mixing, both the ligand and the fluorescently labelled target were centrifuged at 15,000 g, 4°C, for 10 minutes. Maximum concentrations of HMGB1, HMGB1-TL and CXCL12 ligands in the titrations were 104-343 µM, 315 µM and 287-295 µM, respectively. Complex samples were incubated for 30 minutes before loading into NanoTemper premium capillaries. Each experiment was repeated three times, data points are the average of the triplicates and the error bars correspond to the standard deviation.

Since CXCL12 addition induced >10% variation in the fluorescence of 5,6-FAM-Ac-Pep, thermophoresis traces could not be used to measure binding affinity, we therefore used the quenching of the 5,6-FAM-Ac-Pep fluorescence upon binding to estimate the Kd. Data analyses were carried out using NanoTemper Analysis software and the Kd model fitting (one binding site).

### Analytical Ultracentrifugation (AUC)

Sedimentation velocity experiments were performed on an Optima XLI (Beckman Coulter) using an A50 Ti eight-hole rotor and with seven 400 µl samples in standard dual-sector Epon centerpieces equipped with sapphire windows. Absorbance data were acquired at 250 and 280 nm simultaneously with the absorbance scanner in the continuous mode with radial increments of 0.003 cm. Three assembled centrifugation cells containing respectively, free CXCL12 (38.2 µM), HMGB1 (15.6 µM), HMGB1-TL (15.6 µM) and the HMGB1:CXCL12 mixture at the loading ratios of 1:2.5, 1:5, 1:7.5 (with HMGB1 or HMGB1-TL at 7.8 µM concentration), pH 7.5, 20 mM TrisHCl, 50 mM NaCl (or 150 mM NaCl when explicitly stated), were equilibrated at 20°C under vacuum for approximately 1.5 hr prior starting the experiment. Subsequently, centrifugation was performed at 45,000 rpm with 90 scans. The highest protein concentration was determined by the absorbance for which the linear relationship according to Lambert-Beer Law was still guaranteed (O.D. max = 1.0). The buffer density and viscosity were estimated using SEDNTERP^69^.

*c(s) Model*: Sedimentation coefficient distributions at 280 nm of the single proteins were obtained by applying the diffusion-deconvoluted c(s) model, implemented in SEDFIT ^70^. Concentration profiles in terms of absorbance (a(r,t)) were modelled as the sum of Lamm Equation solutions scaled by a continuous distribution c(s) as follows:

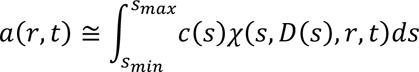

where s is the sedimentation coefficient, (s, D(s), r; t) is the Lamm Equation solution that is dependent on D(s), the corresponding diffusion coefficient, r, radius from the center of rotation, and t, the time from the beginning of the experiment ^32^ .

#### Multisignal Sedimentation Velocity Analysis

Multisignal sedimentation velocity (MS-SV) analysis was performed to determine the stoichiometry of the complex formed by HMGB1 and CXCL12 (or CXCL12-LM). In MS-SV, the standard c(s) approach is modified to deconvolute the contributions of individual species in a component distribution ck(s) where k represents the individual components of a mixture. The absorbance at wavelength λ =(a_λ_, (r, t)) is modelled as:

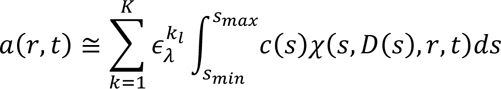

where l is the path length, k is the number of solutes present, and c_k_(s) is a continuous distribution for component k.

MS-SV deconvolution is possible when the complex components have sufficiently different spectral properties, i.e.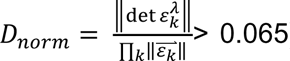)^32^. As the molar extinction coefficients of HMGB1 (ε_280_=20872.8 M^-1^ cm^-1^, ε_250_ = 7987.7 M^-1^ cm^-1^) and CXCL12 (ε_280_=9907.2 M^-1^ cm^-1^, ε_250_=4704.2 M^-1^ cm^-1^) or CXCL12-LM (ε_280_=8413.4 M^-1^ cm^-1^, ε_250_ =4840.5 M^-1^cm^-1^) delivered sufficient spectral discrimination with D_norm_ >0.08 (for HMGB1:CXCL12) and D_norm_ >0.16 (for HMGB1:CXCL12-LM), it was possible to use SEDPHAT to perform global multi-signal analysis of the sedimentation boundary associated to the co-sedimenting complex^71^. MS-SV of the HMGB1:CXCL12 (7.7µM:43µM) and HMGB1:CXCL12-LM (6.9µM:51µM) heterocomplex were collected at 260 nm and 280 nm and globally fit using the multi-wavelength discrete/continuous distribution analysis with mass constraints in SEDPHAT. Integration of the resulting ck(s) distributions revealed the content of each protein component under the peak at a given sedimentation value. Plots of the signal profiles, fits, residuals and the MS-SV results were generated using GUSSI^72^.

### Small Angle X-ray Scattering (SAXS) data collection and analysis

The experiments were performed at the ESRF bioSAXS beamline BM29, Grenoble, France at a detector distance of 2.869 m. CXCL12 and HMGB1 were measured in batch mode at 20°C using the sample changer immediately after protein thawing and centrifugation (30 minutes at 16,000 g). 45 μL of sample solution at three different concentrations (1.95, 3.0 and 3.9 mg/mL per each protein, 20 mM Tris pH 7.5, 50 mM NaCl) were used. To characterize the HMGB1●CXCL12 complex we tested the following protein molar ratio conditions 1:1, 1:2, 1:4 and 1:6 (HMGB1:CXCL12, with HMGB1 1.95 mg/mL). After incubation and centrifugation, the supernatants were immediately measured in the SAXS beamline. Ten frames of 0.5 s/each were collected for each sample (HMGB1●CXCL12, free components at different protein concentration). Data from the HMGB1 and CXCL12 dilutions were merged following standard procedures to create an idealized scattering curve, using PRIMUS within ATSAS 3.1.4^73^. The pair distribution function *p*(*r*) was calculated using GNOM^74^. Protein molecular masses were estimated using both Porod volume and scattering mass contrast methods. All the plots were generated using OriginPro (Version 2022, OriginLab Corporation, Northampton, MA, USA).

To model the HMGB1●CXCL12 heterocomplex, we employed a multistep approach relying on complementary experimental knowledge (NMR and AUC). The SAXS curve of HMGB1 in the presence of two equivalents of CXCL12 was used to generate with EOM a representative starting structure of HMGB1 on which CXCL12 (pdb:2KEE) was docked. Two initial docking models were generated. The first one was obtained with SASREF ^35^ combining the solution scattering data and guiding the docking of CXCL12 (residues 23-28 and 66-67) on HMGB1 (residues 15-45), as suggested by NMR experiments. The second model was obtained docking CXCL12 onto HMGB1 acidic IDR performing a global search with FoXSDock^36^). Next, to describe the dynamic and fuzzy nature of the heterocomplex, we fixed the protein-protein interaction surfaces obtained with SASREF^75^ and FoXSDock^36^, and allowed the rest of HMGB1 to explore a wide range of conformations using the EOM algorithm. The missing residues connecting the rigid bodies and the modelled segments were added with MODELLER^76^. The χ^2^ values of EOM fits over the 0.1-3 nm^-1^ *q*-range of experimental SAXS curves were determined with CORMAP^73^.

All SAXS data were deposited into SASBDB data bank (CXCL12, SASDRG9; HMGB1, SASDRH9; HMGB1●CXCL12, SASDRJ9).

### Molecular images

Molecular images were generated by PyMOL Molecular Graphics System, open source version, Schrödinger, LLC and UCSF Chimera 1.16^77^.

### Cell line and treatments

AB1 mouse malignant mesothelioma cells (MM; Cell Bank, Australia) were cultured in RPMI 1640 (Life Technologies, UK), supplemented with 5% v/v fetal bovine serum (FBS; Life Technologies, UK), 2 mM L-glutamine, and 100 U/ml penicillin/streptomycin at 37°C; 5% CO_2_. Cells were not cultured past passage 10 after cell thawing. To show binding of the HMGB1•CXCL12 heterocomplex to CXCR4, cells were incubated or not at 37°C with increasing concentrations of AMD3100 and then fixed and quantified for the PLA signal. To show the effect of Ac-pep on the binding of the HMGB1•CXCL12 heterocomplex to the receptor, cells were incubated or not at 37°C with increasing concentrations of Ac-pep and then fixed and quantified for the PLA signal.

To measure the binding of the HMGB1•CXCL12 heterocomplex to its receptor, cells were washed with 10 mM Tris-HCl, pH 5.3, washed with RPMI and then incubated for 20 minutes at 4°C with RPMI containing the indicated concentrations of preformed full-length or tailless HMGB1•CXCL12 heterocomplexes. Cells were then fixed and quantified for the PLA signal.

### Proximity ligation assay (PLA)

AB1 cells (2×10^4^) were seeded onto 15 mm glass coverslips and incubated overnight at 37°C/5% CO_2_, and treated the following day as described above . To perform PLA, cells were fixed for 10 minutes in 4% paraformaldehyde in PHEM buffer (1:1) at room temperature (RT), washed in 0.2% bovine serum albumin (BSA)/PBS and blocked with 10% goat serum in 4% BSA/PBS for 1 hr at RT. Cells were then incubated overnight at 4°C with two primary antibodies: mouse monoclonal anti-HMGB1 (1:1000; HMGBiotech) and goat polyclonal anti-CXCL12 (1:50; R&D Systems, #AF-130-NA). Following three washes with 0.2% BSA/PBS, cells were incubated with secondary oligonucleotide-linked antibodies for 1 hr at 37°C (anti-mouse PLUS and anti-goat MINUS, Duolink, Sigma-Aldrich), and processed according to the Duolink PLA fluorescence protocol (Sigma). Finally, cells were stained with Phalloidin-FITC (1:500; Sigma-Aldrich, # P5282) and Hoechst (1 μg/mL; Sigma-Aldrich, #33342), and mounted on microscope slides with Flourosave reagent (Merck, #345789).

All images for PLA were acquired using a Leica TCS SP5 X confocal microscope (Leica Biosystems) with a 63x objective, using channels for Hoechst (405 nm), FITC (488 nm), and Texas Red (561 nm). Z-step size was set at 0.69 µm and the top and bottom of the cells were ascertained manually prior to acquiring each image. Each image was stacked to max intensity using ImageJ/Fiji software and saved as tiff files. These images were then used to create cytoplasmic masks using Cellpose^78, 79^. Python 3.10.0 was used to run Cellpose. Cytoplasmic masks were used to ascertain cytoplasmic specific PLA signal in ImageJ/Fiji. Data are expressed as RawIntegratedDensity (sum of all pixel intensities in region of interest, in arbitrary units) normalized to the area of the respective cytosolic regions.

## Author contributions

MVM produced recombinant proteins. MVM; FDL, MG and GQ analyzed NMR experiments. MVM, TS and SR performed and analyzed ITC experiments. MVM performed and analyzed AUC experiments and prepared a draft of the manuscript. GQ performed the NMR experiments. FDL and CZ performed and analyzed MST experiments. LC, FDM and MC performed and analyzed PLA experiments. GG, MVM and FDL performed SAXS experiments at ESRF BM29; GG and MG analyzed SAXS experiments. MEB supervised the cell-based experiments. GM supervised the study and was involved in all aspects of the experimental design, data analysis, and manuscript preparation. All authors critically reviewed the text and figures.

## Conflict of interest

The authors declare that they have no conflict of interest. However, LC is an employee and MEB is founder and part-owner of HMGBiotech, a company that provides goods and services related to HMGB proteins.

## Acknowledgements

The research leading to these results has received funding from AIRC under IG 2018 - ID. 21440 project – P.I. Giovanna Musco

## Supporting Information

Supplementary Figures

Supplementary Tables

