## Supplementary Figures and Tables for "The acidic intrinsically disordered region of the inflammatory mediator HMGB1 mediates fuzzy interactions with chemokine CXCL12"

### Supplementary Information

**Supplementary Figures:** page 2-6

**Supplementary Tables:** page 7-9

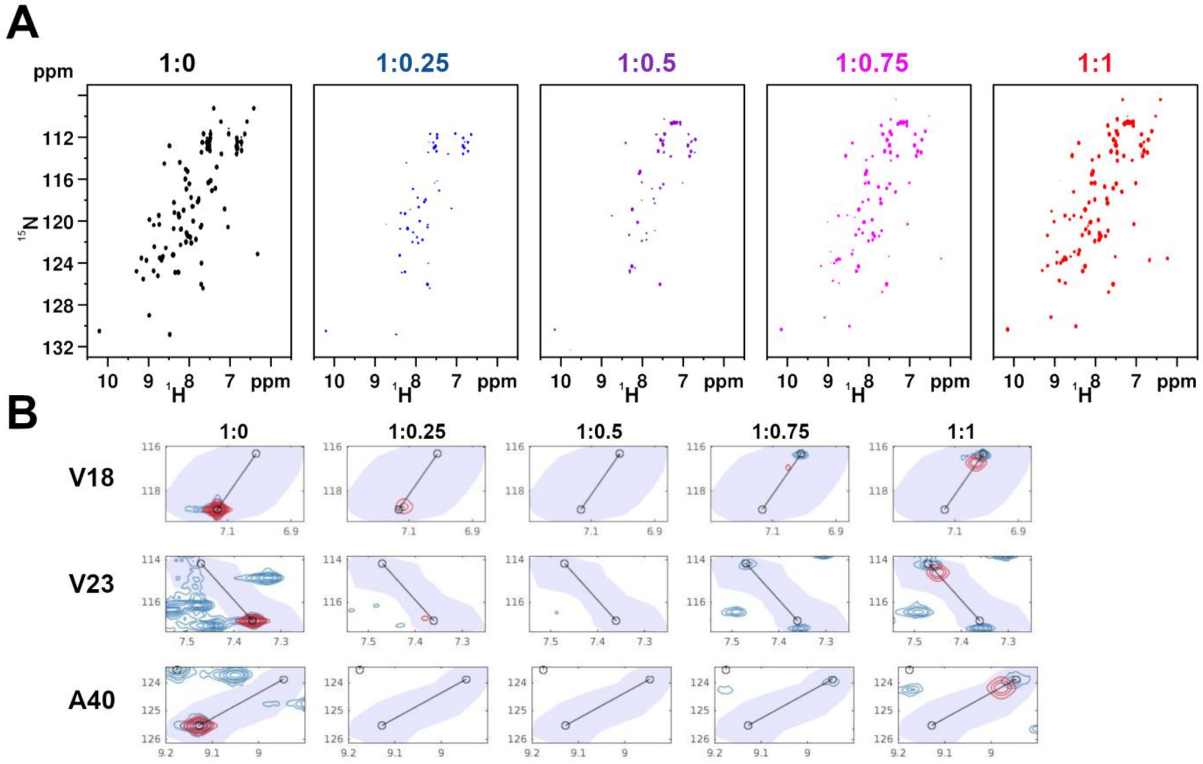


**Supplementary Figure S1: Binding of Ac-Pep to CXCL12 (A)** $^{1}$H-$^{15}$N HSQC spectra of $^{15}$N CXCL12 (0.1 mM) in the presence of increasing concentrations of Ac-pep. Peaks disappear at sub-stoichiometric ratio and reappear at equimolar ratio. **(B)** TITAN fits using single-site binding equation for CXCL12 resonance-specific trajectories as a function of Ac-pep. Original spectra are shown in blue contours, fitted spectra are shown in red contours.

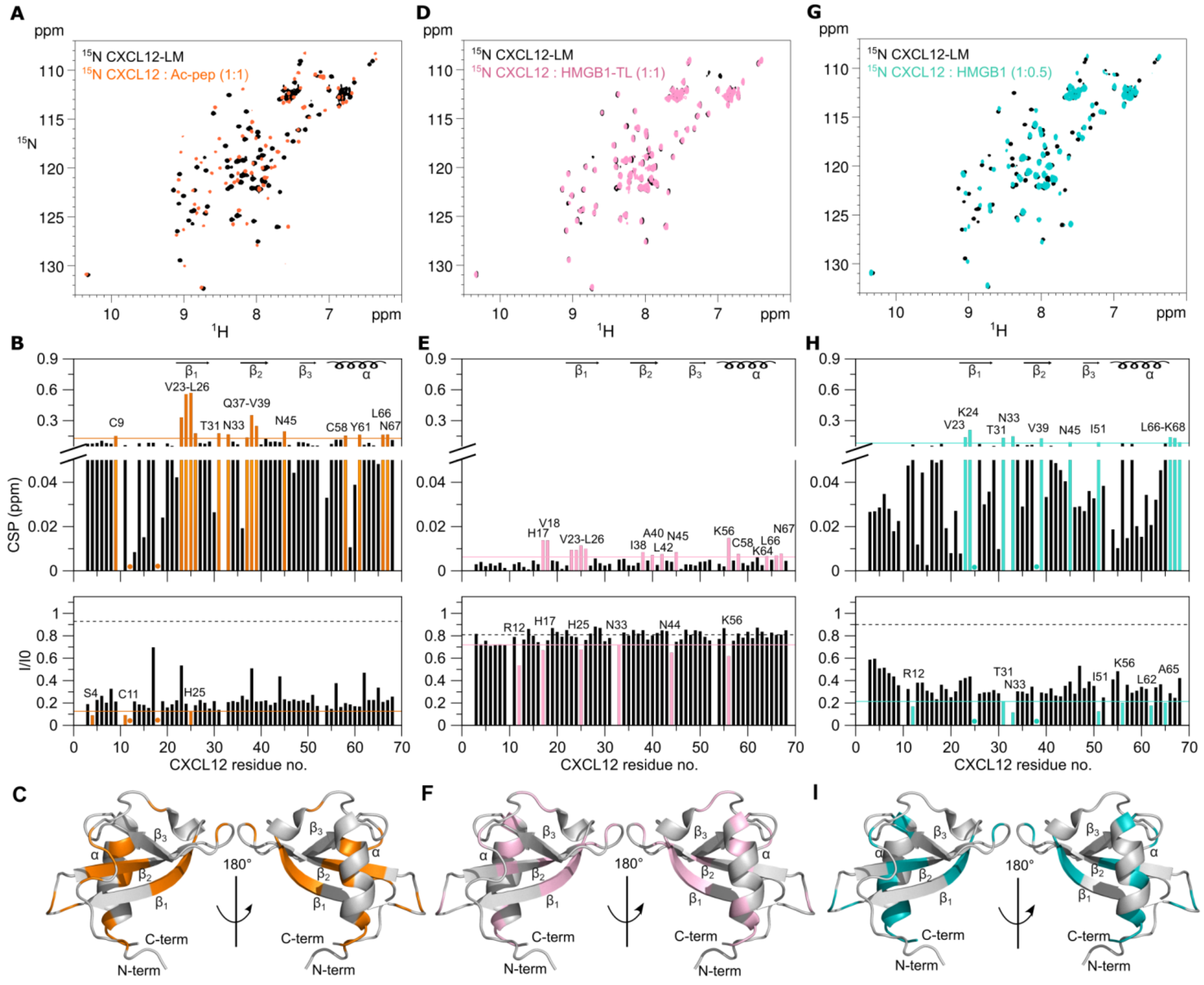


**Supplementary Figure S2: The interaction of CXCL12-LM with Ac-pep, HMGB1-TL and HMGB1 is similar to CXCL12 (A)** Superposition of $^{1}$H-$^{15}$N HSQC spectra of $^{15}$N CXCL12-LM (0.1 mM) without (black) and with (orange) Ac-pep (1:1). **(B)** *Upper panel*: Bar graph showing residue-specific CSPs of $^{15}$N-labeled CXCL12-LM (0.1 mM) upon addition of Ac-pep (1:1). Residues with CSP > Avg + $\sigma_0$ (orange line) are labeled and represented in orange. *Lower panel***:** Bar graph showing residue-specific peak intensities ratios ($I/I_0$) of $^{15}$N-labeled CXCL12-LM (0.1 mM) upon addition of Ac-pep (1:1). Residues with $I/I_0$ < Avg – SD (orange line) are shown in orange. (**C**) CXCL12-LM (gray cartoon, pdb code: 2n55) residues with CSP > Avg + $\sigma_0$ and with $I/I_0$ < Avg - SD are shown in orange **(D)** Superposition of $^{1}$H-$^{15}$N HSQC spectra of $^{15}$N CXCL12-LM (0.1 mM) without (black) and with (pink) HMGB1-TL (1:1). **(E)** *Top panel*: Bar graph showing residue-specific CSPs of $^{15}$N-labeled CXCL12-LM (0.1 mM) upon addition of HMGB1-TL (1:1). Residues with CSP > Avg + $\sigma_0$ (pink line) are represented in pink and labeled. *Lower panel*: Bar graph showing residue-specific peak intensities ratios ($I/I_0$) of $^{15}$N-labeled CXCL12-LM (0.1 mM) upon addition of Ac-pep (1:1). Residues with $I/I_0$ < Avg - SD (pink line) are shown in pink **(F)** CXCL12-LM (gray cartoon, pdb code:2n55). CXCL12-LM residues with CSP > Avg + $\sigma_0$ and with $I/I_0$< Avg - SD are shown in purple. **(G)** Superposition of $^{1}$H-$^{15}$N HSQC spectra of $^{15}$N CXCL12-LM (0.1 mM) without (black) and with (cyan) HMGB1 (1:1). **(H)** *Top Panel*: Bar graph showing residue-specific CSPs of $^{15}$N-labeled CXCL12-LM (0.1 mM) upon addition of HMGB1 (1:1). Residues with CSP > Avg + $\sigma_0$ (cyan line) are represented in green and labeled. *Lower panel*: Bar graph showing residue-specific peak intensities ratios ($I/I_0$) of $^{15}$N-labeled CXCL12-LM (0.1 mM) upon addition of Ac-pep (1:1). Residues with $I/I_0$ < Avg - SD (cyan line) are shown in green. (**I**) CXCL12-LM (gray cartoon, pdb code:2n55). CXCL12-LM residues with CSP > Avg + $\sigma_0$ (cyan line) and with $I/I_0$ < Avg - SD are shown in cyan. In the bar-graphs α-helices and β-strands are schematically represented on the top, missing residues are prolines or are absent because of exchange with the solvent, dots indicate resonances disappeared upon binding; the dashed black line indicates the peak intensity decrease due to the titration dilution effect.

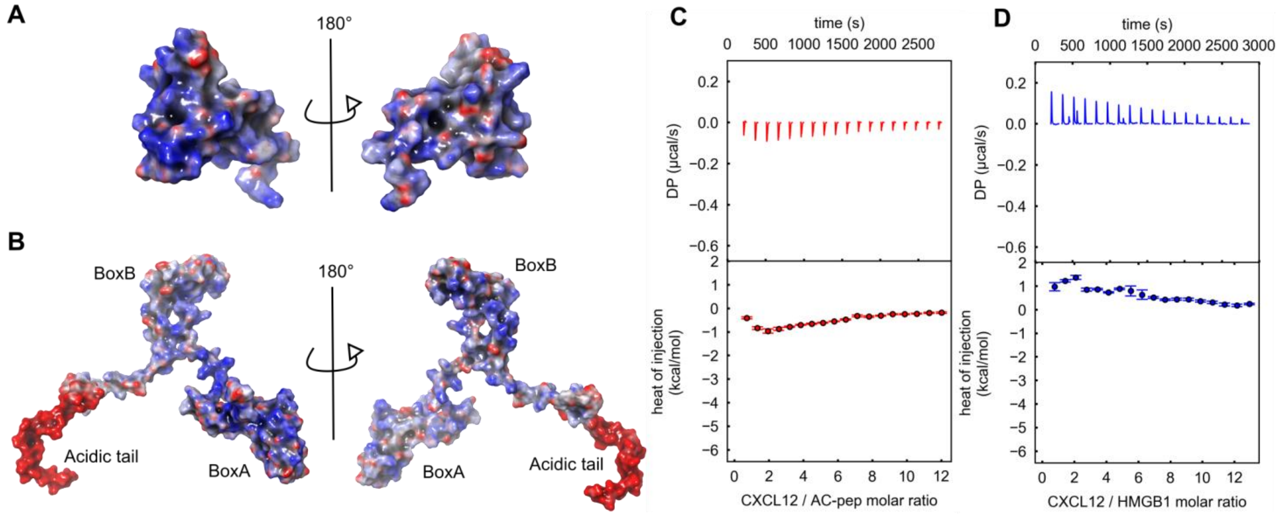


**Supplementary Figure S3: Electrostatic interactions drive enthalpic changes in HMGB1•CXCL12 complex formation.** Electrostatic surface representation (as calculated by the Poisson Boltzmann Electrostatic Surface routine in Maestro-Schroedinger) of **(A)** CXCL12 and **(B)** HMGB1. The ITC sequential heat pulses (upper panel) and the integrated data corrected for heat of dilution (lower panel) of CXCL12 binding to **(C)** Ac-pep (20 mM TrisHCl at pH 7.5, 150 mM NaCl) and to **(D)** HMGB1 (20 mM TrisHCl at pH 7.5, 150 mM NaCl).

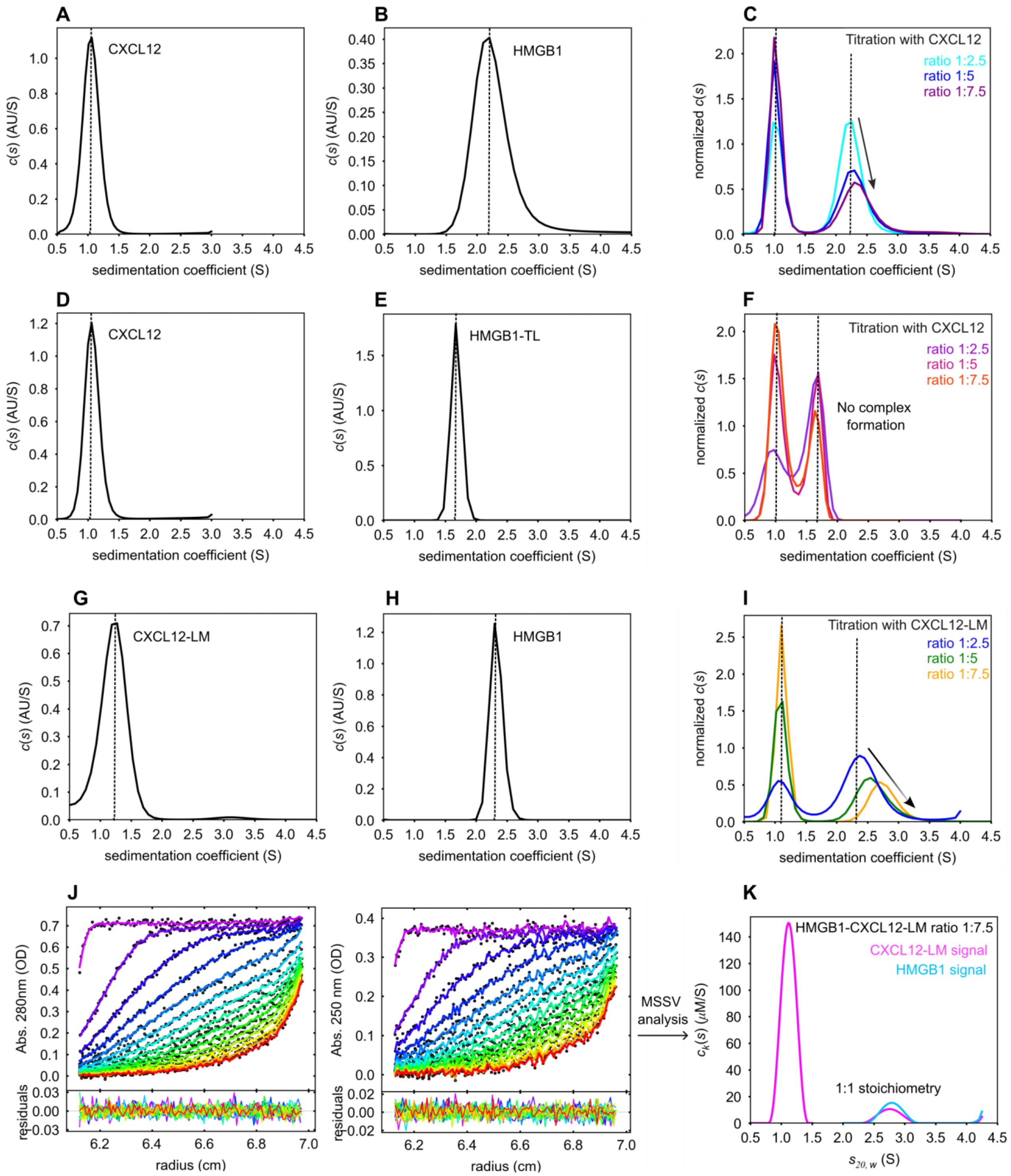


**Supplementary Figure S4: Analytical ultracentrifugation analysis of the interaction between: (A-C) CXCL12 and HMGB1 (150 mM NaCl), (D-F) CXCL12 and HMGB1-TL (50 mM NaCl) and (G-K) CXCL12-LM and HMGB1 (50 mM NaCl)** Sedimentation velocity analytical ultracentrifugation (SV-AUC) experiments of free (**A**) CXCL12 and (**B**) HMGB1, scanned by absorbance at 280nm. (**C**) SV-AUC analysis of the interaction between HMGB1 (7.8 µM) and increasing concentrations of CXCL12 (colors). To facilitate comparison the dotted lines indicate the sedimentation coefficients of the free components. The continuous sedimentation coefficient distribution analyses (c(s)) shows that the peaks corresponding to HMGB1 broadens and shifts to a larger s-value in an CXCL12 concentration-dependent manner. Experimental conditions: pH 7.5, 20 mM TrisHCl, 150 mM NaCl. Sedimentation velocity analytical ultracentrifugation (SV-AUC) experiments of free (**D**) CXCL12 and (**E**) HMGB1-TL, scanned by absorbance at 280nm. (**F**) The sedimentation coefficient distribution analyses (c(s)) of the complex shows that the peak corresponding to HMGB1 does not shift to larger c(s)-values in a CXCL12 concentration-dependent manner, suggesting that the association between the two components is too rapid to be detected in the sedimentation time scale. The dotted lines indicate the sedimentation coefficients of the free components. Experimental conditions: pH 7.5, 20 mM TrisHCl, 50 mM NaCl. Sedimentation velocity analytical ultracentrifugation (SV-AUC) experiments of free (**G**) CXCL12-LM and (**H**) HMGB1, scanned by absorbance at 280nm. (**I**) Analysis of the interaction between HMGB1 (7.8 µM) and increasing concentrations of CXCL12-LM (colors). To facilitate comparison the dotted lines indicate the sedimentation coefficients of the free components. The sedimentation coefficient distribution analyses (c(s)) shows that the peaks corresponding to HMGB1 broadens and shifts to a larger s-value in an CXCL12-LM concentration-dependent manner. (**J**) Global multi-signal sedimentation velocity (MS-SV) analysis to determine the stoichiometry of HMGB1:CXCL12-LM complex, with 6.9µM HMGB1 and 51 µM CXCL12. The raw sedimentation signals of the HMGB1●CXCL12-LM heterocomplex acquired at different time points with absorbance at 280 nm (left), and absorbance at 250 nm (right) with the signal profiles shown in units OD280, and OD250, respectively, as a function of radius in centimeters. The time-points of the boundaries are indicated in rainbow colors, progressing from purple (early scans) to red (late scans). Only every 3rd data point used in the analysis are shown. Residuals of the fit are shown at the bottom **(K).** Global multi-wavelength analysis and decomposition into the component sedimentation coefficient distributions, ck(s), for CXCL12-LM (magenta) and HMGB1 (cyan line). Integration of sedimentation coefficient distributions of CXCL12-LM and HMGB1 yields the concentration of the co-sedimenting components in the reaction mixture (**Supplementary Table S1**).

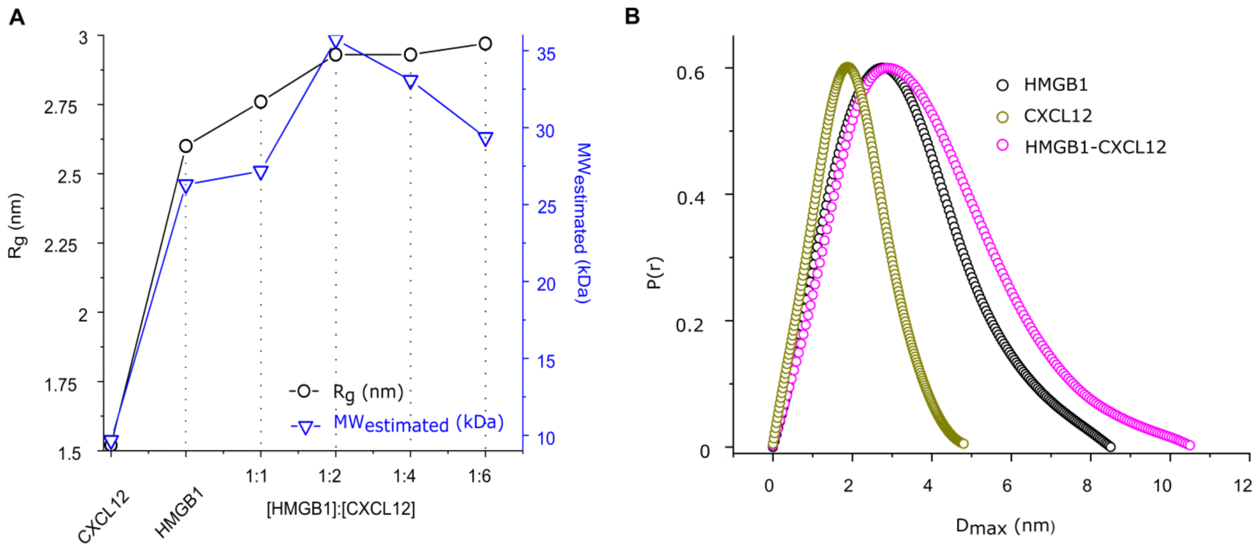


**Supplementary Figure S5: Structural parameters of the HMGB1●CXCL12 complex. (A)** Titration of HMGB1 with increasing CXCL12 equivalents and estimation of the $R_g$ (in black) and molecular weight (MW, in blue) parameters of the assemblies. At 1:4 and 1:6 conditions a decrease in MW estimation is visible, due to the contribution of free CXCL12. The $R_g$ and MW values are stable at 1:2, suggesting that in this condition the HMGB1●CXCL12 complex is stable. **(B)** Distance probability plot with the *P*(*r*) versus $D_{max}$ (in nm) profiles from the SAXS experimental data of CXCL12 (gold circles), HMGB1 (black circles) and HMGB1-CXCL12 complex (magenta circles).

### Supplementary Table S1: Calculated values from AUC analysis of CXCL12-LM, HMGB1 and HMGB1●CXCL12-LM, pH 7.5, 20 mM TrisHCl, 50 mM NaCl.

| | *C(s) analysis of single components* | | | *MS-SV model analysis HMGB1:CXCL12-LM 1:7.5* | | |
|---|---|---|---|---|---|---|
| | *Param.* | **CXCL12-LM** | **HMGB1** | *Param.* | **CXCL12-LM$_{mix}$** | **HMGB1$_{mix}$** |
| AUC-SV | *sw*[1] | 1.2 S | 1.7 S | $sw^{5}_{(20,w)\ complex}$ | 2.8 S | 2.9 S |
| | $f/f_0^{2}$ | 1.2 (globular) | 1.6 (elongated) | $conc^{6}_{tot}$ | 51 µM | 6.9 µM |
| | *MW (kDa)*[3] | 8.9 | 21.1 | $conc^{7}_{complex}$ | 4.9 µM | 6.5 µM |
| | *RMSD*[4] | 0.0057 | 0.0049 | $RMSD^{8}_{280nm}$<br>$RMSD^{9}_{250nm}$<br>$D_{norm}^{10}$ | 0.0077<br>0.0057<br>0.16 | |

[1]sedimentation coefficient
[2]frictional ratio, f is the frictional parameter of the particle and $f_0$ is the frictional value of a smooth sphere
[3]estimated molecular weight
[4]RMSD, root mean square deviation of the fitting
[5]sedimentation coefficient normalized to standard solution conditions of water at 20° C expressed in Svedberg (S)
[6]concentration of the components; $conc_{tot}$
[7]calculated concentration of the components in the complex; $conc_{complex}$
[8]root mean square deviation of the fitting at 280 nm
[9]root mean square deviation of the fitting at 250 nm
[10]$D_{norm}$ determinant of the extinction coefficient matrix.

### Supplementary Table S2: Calculated values from AUC analysis of CXCL12, HMGB1 and HMGB1●CXCL12, pH 7.5, 20 mM TrisHCl,150 mM NaCl.

| | *C(s) analysis of single components* | | | *MS-SV model analysis HMGB1:CXCL12 1:7.5* | | |
|---|---|---|---|---|---|---|
| | *Param.* | **CXCL12** | **HMGB1** | *Param.* | **CXCL12$_{mix}$** | **HMGB1$_{mix}$** |
| AUC-SV | *sw*[1] | 1.05 S | 2.3 S | $sw^{5}_{(20,w)\ complex}$ | 2.5 S | 2.54 S |
| | $f/f_0^{2}$ | 1.3 (globular) | 1.7 (elongated) | $conc^{6}_{tot}$ | 48 µM | 7.8 µM |
| | *MW (kDa)*[3] | 8.6 | 34.4 | $conc^{7}_{complex}$ | 6.0 µM | 7.6 µM |
| | *RMSD*[4] | 0.0056 | 0.0077 | $RMSD^{8}_{280nm}$<br>$RMSD^{9}_{250nm}$<br>$D_{norm}^{10}$ | 0.0079,<br>0.0058<br>0.09 | |

[1]sedimentation coefficient
[2]frictional ratio, f is the frictional parameter of the particle and $f_0$ is the frictional value of a smooth sphere
[3]estimated molecular weight
[4]RMSD, root mean square deviation of the fitting

### Supplementary Table S3. Experimental details on SAXS experiments and analysis

*(A) Sample details*

| Sample name | **CXCL12** | **HMGB1** | **HMGB1-CXCL12** |
|---|---|---|---|
| Organism | *Homo sapiens sapiens* | *Rattus norvegicus* | - |
| UniProt sequence ID | P48061 | P63159 | - |
| Calculated molecular weight (Da) | 9183.88 | 24748.54 | 33932.42 |
| Total frames (frames used) | 10 (8) | | |
| Protein concentration (mg/mL) | 1.95, 3.0, 3.9 | | - |
| Molar ratii (HMGB1:CXCL12) | | | 1:1, 1:2, 1:4, 1:6 |
| Protetin buffer | 20 mM Tris pH 7.5, 50 mM NaCl | | |

*(B) SAXS data collection parameters*

| | |
|---|---|
| Instrument | ESRF BM29 |
| Wavelength (Å) | 0.99 |
| *q*-range (Å) | 0.004-0.5 |
| Sample-to-detector distance (m) | 2.867 |
| Exposure time | 0.5 sec/frame |
| Temperature (° C) | 20 |
| Detector | Pilatus3 X 2M (Dectris) |
| Flux (photons/s) | $2 \times 10^{12}$ |
| Beam size (μm) | 100 x 100 |
| Sample configuration | 1.8 mm quartz glass capillary |
| Absolute scaling method | Comparison to water in sample capillary |
| Normalization | To transmitted intensity by beam-stop counter |

*(C) Structural parameters*

| | **CXCL12** | **HMGB1** | **HMGB1-CXCL12** |
|---|---|---|---|
| **Guinier analysis** | | | |
| I(0) | 32.64 | 43.98 | 82.69 |
| $R_g$ (nm) | 1.52 ± 0.006 | 2.6 ± 0.018 | 2.93 ± 0.016 |
| $q$-range ($nm^{-1}$), point range | 0.035-0.746, 20-140 | 0.0343-0.27, 13-73 | 0.0243-0.22, 10-71 |
| ***P*(r) analysis** | | | |
| I(0) ($cm^{-1}$) | 32.8 | 43.84 | 83.1 |
| $R_g$ (nm) | 1.54 | 2.62 | 3.0 |
| $D_{max}$ ($R_{max,}$ nm) | 4.8 | 8.5 | 10.5 |
| $q$-range ($nm^{-1}$), point range | 0.035-4.0, 11-752 | 0.156-3, 13-573 | 0.0332-2.8, 7-508 |
| Porod volume ($nm^3$) | 19.352 | 52.729 | 71.738 |
| $\chi^2$ [total estimate from GNOM] | 0.8897 | 0.8873 | 0.8573 |
| Mass estimate based on volume (kDa), ratio to predicted | 9.676, 1.05 | 26.36, 1.06 | 35.82, 1.05 |
| Mass estimate based on Bayesian inference (kDa), ratio to predicted | 9.5, 1.03 | 28.23, 1.1 | 38.834, 1.1 |

*(D) Software employed for SAXS data reduction, analysis and interpretation*

| | |
|---|---|
| SAXS data reduction and processing | EDNA and Primus (ATSAS 3.2.1) |
| Shape/bead modelling | DAMMIF(ATSAS 3.2.1) |
| Atomic structure modelling | EOM (ATSAS 3.2.1) |
| 3D graphic representation | UCSF Chimera 1.15 |
| Missing sequence modelling | MODELLER |
| SAXS-based docking | SASREF (ATSAS 3.2.1), FoXSDock (version main.ec6dbc2) |

*(E) Shape model-fitting results and rigid-body atomistic modelling*

| | **CXCL12** | **HMGB1** | **HMGB1-CXCL12** | |
|---|---|---|---|---|
| **EOM** | *Default parameters, 10,000 models in initial ensemble, native-like models, constant subtraction allowed* | | | |
| | | | **EOM-SASREF** | **EOM-FoXSdock** |
| PDB id for rigid bodies | - | - | 2KEE (CXCL12), Alphafold P63159 (HMGB1) | |
| Rigid bodies (residue numbers) | - | - | 8-68 (CXCL12); 13-76 (BoxA); 99-164 (BoxB) | 8-68 (CXCL12); 13-76 (BoxA); 99-164 |

| | | | | |
|---|---|---|---|---|
| | | | | (BoxB); 187-205 (acidic IDR) |
| Position of rigid bodies in the space during simulation | - | - | CXCL12 (fixed); BoxA (fixed); BoxB (free) | CXCL12 (fixed); BoxA (free); BoxB (free); acidic IDR (fixed) |
| Segments modelled *via* EOM (residue numbers) | - | - | 1-7 (CXCL12); 1-12, 77-98, 165-214 (HMGB1) | 1-7 (CXCL12); 1-12, 77-98, 165-186; 206-214 (HMGB1) |
| $\chi^2$ value for the fitting | - | - | 0.975 | 0.979 |
| CORMAP *P*-values | - | - | 0.422 | 0.422 |
| Number of representative structures | - | - | 5 | 4 |
| $R_{flex}$ ensemble (%) | - | - | 69.38 | 69.16 |
| $R_{flex}$ pool (%) | - | - | 84.27 | 85.04 |
| $R_{\sigma}$ ensemble | - | - | 0.63 | 0.72 |
| Weighted average $R_g$ (nm) | - | - | 2.99 | 2.95 |
| Weighted average $D_{max}$ (nm) | - | - | 9.19 | 9.88 |
| % fraction of representative structures (model number) | - | - | 10 % (1), 5 % (2), 5 % (3), 65 % (4), 15 % (5) | 20 % (1), 70 % (2), 5 % (3), 5 % (4) |
| **SASREF** | *Default parameters* | | | |
| PDB id for rigid bodies | - | | 2KEE (CXCL12), Alphafold P63159 (HMGB1 | |
| Contact region; max distance (Å) | - | - | Residues 23-28 (CXCL12), 34-40 (HMGB1); 5 Å | |
| $\chi^2$ value for the fitting | - | - | 1.2 | |
| **FoXSDock** | *Default parameters* | | | |
| PDB id for rigid bodies | - | - | 2KEE (CXCL12), Alphafold P63159 (HMGB1 | |
| Contact region; max distance (Å) | - | - | Residues 15-28 (CXCL12), 170-214 (HMGB1); 5 Å | |
| $\chi^2$ value for the fitting | - | - | 4.2 | |

*(F) Small Angle Scattering Biological Data Bank (SASBDB)*

| | CXCL12 | HMGB1 | HMGB1-CXCL12 |
|---|---|---|---|
| SASBDB ID* | SASDRG9 | SASDRH9 | SASDRJ9 |
| *: https://www.sasbdb.org/project/2028/plawdyohws/ | | | |
